# Rv3722c governs aspartate-dependent nitrogen metabolism in *Mycobacterium tuberculosis*

**DOI:** 10.1101/784462

**Authors:** Robert Jansen, Lungelo Mandyoli, Ryan Hughes, Shoko Wakabayashi, Jessica Pinkham, Bruna Selbach, Kristine Guinn, Eric Rubin, James Sacchettini, Kyu Rhee

**Affiliations:** Division of Infectious Diseases, Department of Medicine, Weill Cornell Medical College, New York, New York 10065, USA; Department of Biochemistry and Biophysics, Texas A&M University, College Station, TX; Department of Immunology and Infectious Diseases, Harvard T. H. Chan School of Public Health, Boston, Massachusetts, USA; Department of Microbiology & Immunology, Weill Cornell Medical College, New York, New York 10065, USA

## Abstract

Organisms are defined by their genomes, yet many distinguishing features of a given organism are encoded by genes that are functionally unannotated. *Mycobacterium tuberculosis* (*Mtb*), the leading cause of death due to a single microbe, co-evolved with humans as its only known natural reservoir, yet the factors mediating *Mtb’s* pathogenicity remain incompletely defined. *rv3722c* is a gene of unknown function predicted to encode a pyridoxal phosphate binding protein and to be essential for *in vitro* growth of *Mtb*. Using metabolomic, genetic and structural approaches, we show that Rv3722c is the primary aspartate aminotransferase of *Mtb* and mediates an essential but underrecognized role in metabolism: nitrogen distribution. Together with the attenuation of Rv3722c-deficient *Mtb* in macrophages and mice, these results identify aspartate biosynthesis and nitrogen distribution as potential species-selective drug targets in *Mtb*.

## Introduction

The accumulation of sequence data is outpacing the ability to annotate its information content^1–3^. As many as half of all predicted coding sequences are estimated to be unannotated or misannotated or to encode additional activities beyond their predicted functions^1, 4^. Conversely, approximately one third of all detected enzymatic activities lack an associated coding sequence^5^. For microbial pathogens, such gaps have restricted access to what may be the most specific and actionable features of their physiology.

*Mycobacterium tuberculosis* (*Mtb*) is the causative agent of tuberculosis (TB) and the leading cause of death due to an infectious agent^6^. Curative chemotherapies for TB were first developed over 50 years ago but are only recently beginning to change and currently number among the longest, most complex and toxic treatments for a bacterial infection. Together, these shortcomings have fostered rates of treatment non-compliance and default that help fuel the pandemic and promote the emergence of drug resistance. Shorter, simpler chemotherapies are urgently needed.

*rv3722c* is an *Mtb* gene of unknown function, predicted by transposon mutagenesis to be essential for *in vitro* growth^7, 8^. Bio-informatic analyses predict that *rv3722c* encodes a pyridoxal phosphate (PLP) binding domain. Consistent with that prediction, Rv3722c is annotated as an aminotransferase^9, 10^, but it is also annotated as a member of the GntR family of transcription factors^11, 12^, while experimental studies have implicated Rv3722c as a serine hydrolase^13^ and a secreted protein^14, 15^.

Growing evidence has implicated carbon metabolism as a determinant of *Mtb* pathogenicity^16^. However, recent work has begun to implicate a similarly important role for nitrogen uptake and assimilation^17–19^. Here, we demonstrate that Rv3722c encodes the primary aspartate aminotransferase (AspAT) in *Mtb,* non-redundantly catalyzes the specific biosynthesis of Asp *in vitro* and is essential for axenic growth and survival of *Mtb* in macrophages and in mice. We further show that this essentiality is due, in part, to a non-redundant role of Asp in the metabolic distribution of assimilated nitrogen. These findings identify Rv3722c as an essential metabolic mediator of Asp biosynthesis and Asp-dependent nitrogen metabolism as an essential determinant of *Mtb* growth and virulence.

## Results

### Rv3722c is conditionally essential *in vitro*

We first sought to confirm the predicted essentiality of Rv3722c. We constructed an *Mtb* strain in which expression of Rv3722c is regulated by its native promoter, but protein stability is controlled by a tetracycline-repressible protein degradation system (Rv3722c-TetON)^20, 21^. An isogenic strain lacking a functional protein degradation system (Rv3722c-Control) served as a control. As predicted, omission of anhydrotetracycline (ATC) resulted in depletion of Rv3722c protein and attenuation of *Mtb* growth in Glu-based Middlebrook 7H9 medium (Fig 1A-B). This growth defect could be overcome by culturing Rv3722c-TetON in the same medium supplemented with casein hydrolysate, which contains a complex mixture of amino acids and small peptides (Fig 1C), and mitigated by culture on Middlebrook 7H10 agar (Fig 1D). This conditional rescue made it possible to deplete Rv3722c to levels below the limit of detection (Fig 1E) before a subsequent experimental challenge. Such pre-depletion completely prevented subsequent growth in unsupplemented 7H9 (Fig 1F). Rv3722c could also be depleted below the limit of detection without impairing growth by culturing the cells in an Asn-based minimal medium (Sauton’s) (Fig S1).

**Figure 1.**
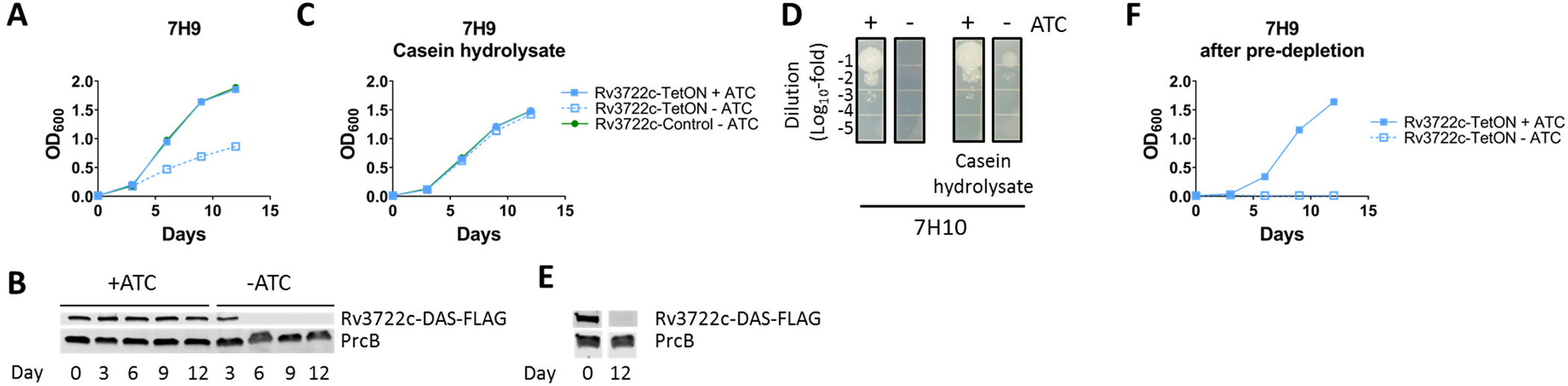
Rv3722c is conditionally esssential *in vitro*. **A)** Growth curve of Rv3722c-proficient and – deficient *Mtb* in 7H9 culture media. Rv3722c-TetOn and Rv3722c-control were cultured in Middlebrook 7H9 culture media with or without 500 ng/mL anhydrotetracycline (ATC). Bacterial growth was monitored by optical density at 600 nm. **B)** Western blot showing the depletion of Rv3722c 7H9 culture media. Rv3722c-TetON was cultured in 7H9 with or without ATC for 12 days (corresponding to A). Protein lysates were analyzed by Western blotting, using an α-FLAG antibody. The proteasome subunit β (PrcB) was used as loading control. **C)** Growth curve of Rv3722c-proficient and –deficient *Mtb* in 7H9 supplemented with casein hydrolysate. As A, but using 7H9 supplemented with 1% casein hydrolysate. **D)** Spot assay of Rv3722c-proficient and –deficient *Mtb* on solid growth media. A serially diluted Rv3722c-TetON culture (OD 0.1) was spotted onto Middlebrook 7H10 agar with or without 1% casein hydrolysate, in the presence or absence of ATC, and cultures for 2 weeks. **E)** Western blot showing depletion of Rv3722c. As B, but in 7H9 supplemented with 1% casein hydrolysate. **F)** Growth curve of of Rv3722c-proficient and –pre-depleted *Mtb* in 7H9 growth media. Rv3722c-TetOn in 7H9 with or without ATC, after pre-depletion of Rv3722c in 7H9 with 1% casein hydrolysate without ATC. For all growth curves, data are represented as mean -/+ SD of three experimental replicates (n=3) representative of at least two independent experiments. See also Fig S1.

### Rv3722c is essential for infection

We next tested the essentiality of Rv3722c *in vivo* by first infecting mouse bone marrow-derived macrophages with Rv3722c-sufficient and Rv3722c-pre-depleted *Mtb*. Growth of Rv3722c-deficient *Mtb* was severely compromised in both resting and interferon-γ-activated macrophages (Fig 2A). Testing in an aerosol infection model of TB in mice similarly revealed a striking attenuation of Rv3722c-deficient *Mtb* in lungs, as reported by a lack of growth (Fig 2B) or their rapid and complete replacement by escape mutants that were no longer ATC-responsive (Fig S2).

**Figure 2.**
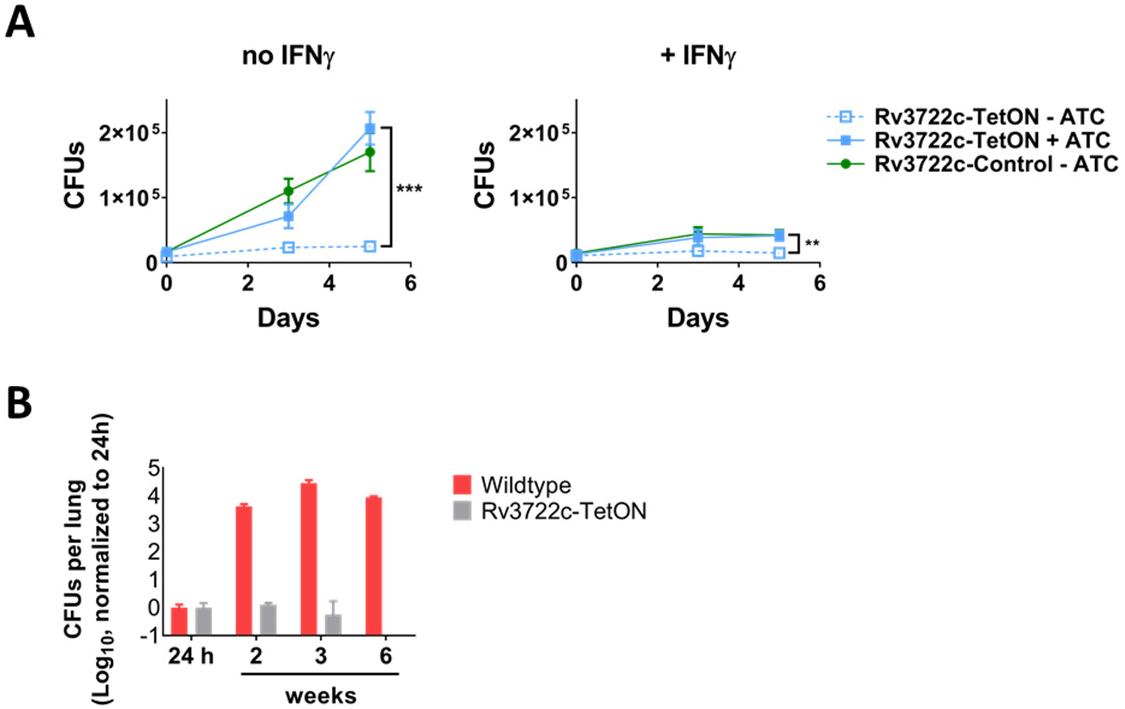
Rv3722c is required for virulence in macrophages and mice. **A)** Timecourse of macrophage infection. Primary murine bone marrow-derived macrophages were treated with control or interferon γ (IFNγ) for activation, followed by infection with Rv3722c-control and Rv3722c-TetON precultured in 7H9+casein hydrolysate with or without ATC to pre-deplete Rv3722c (multiplicity of infection of 1). The number of colony-forming units (CFU’s) were assessed by plating serial dilutions on 7H10 solid media supplemented with casein hydrolysate and ATC. Data are presented as mean +/- SD of three experimental replicates (n=3) representative of two independent experiments; **: p< 0.01, ***:p<0.001 using unpaired Student’s *t*-test. **B)** Timecourse of mouse infection. Mice were infected with aerosolized wildtype *Mtb* and Rv3722c-TetON pre-cultured in Sauton’s without ATC to generate Rv3722c-deficient *Mtb*. After infection, mice were fed chow without doxycycline. The number of CFU’s were assessed by plating serial dilutions on 7H10 solid media with and without ATC to determine the number of non-ATC-responsive mutants. Because the Rv3722c-TetON was lower that that of wildtype, the number of CFUs were normalized to the number of CFU’s detected 24h. No non ATC-dependent Rv3722c-TetON mutants were detected. Data are represented as mean +/- SD (n=3-5). A similar experiment is shown in Fig S2.

### Rv3722c functions as an aminotransferase

Based on bio-informatic evidence implicating Rv3722c as a PLP-dependent protein, we assayed purified recombinant Rv3722c for catalytic activity using activity-based metabolomic profiling (ABMP). ABMP is a biochemically unbiased method developed to ‘de-orphan’ unannotated metabolic enzymes that consists of incubation of a protein of interest with a highly concentrated homologous metabolite extract and monitoring for time- and protein-dependent changes indicative of catalysis by untargeted high-resolution mass spectrometry^22–26^. ABMP with Rv3722c identified several putative product features (fragments, adducts, dimers and isotopes) that could all be related to a metabolite with m/z 144.03 [M-H]^-^ (Fig 3A). This metabolite was formed in a time- and Rv3722c-dependent fashion (Fig 3B), and identified by mass spectral fragmentation as ketoglutaramate (keto-Gln), the ketoacid of Gln (Fig 3C).

**Figure 3.**
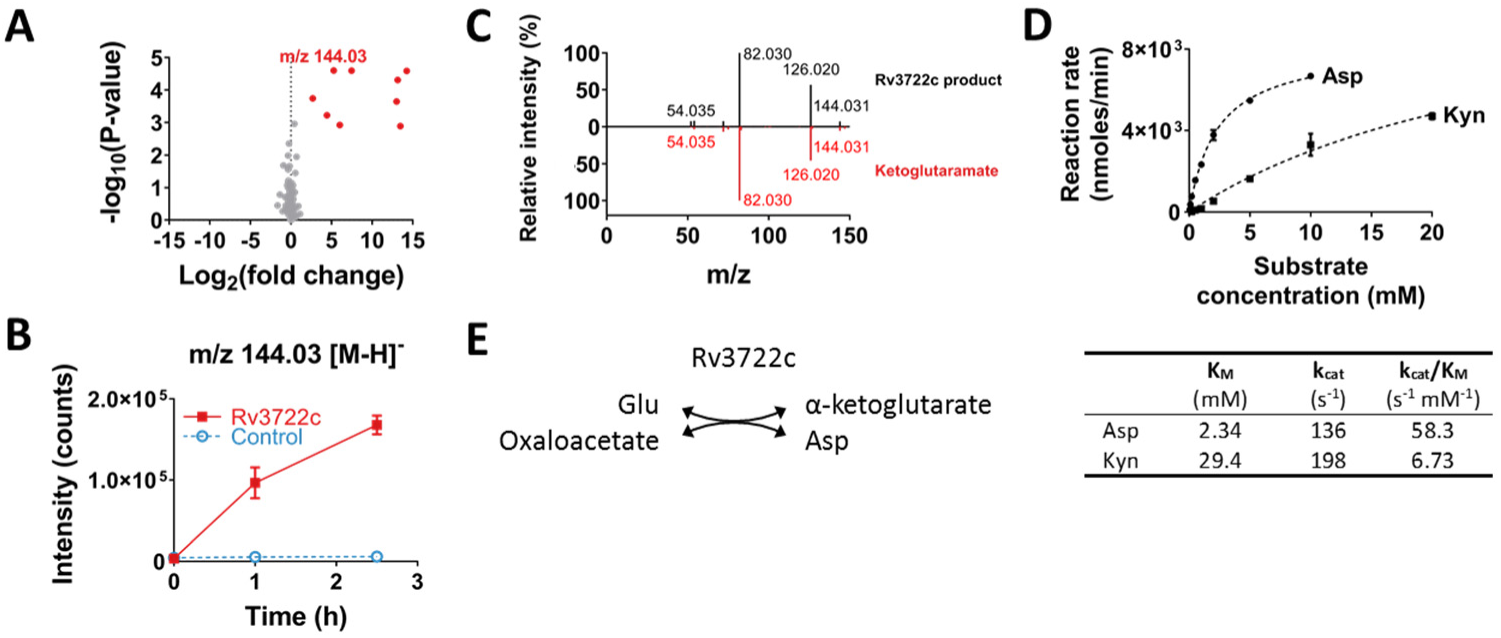
Rv3722c functions as an aspartate aminotransferase. **A)** Volcano-plot of activity-based metabolite profiling with purified recombinant Rv3722c. Purified recombinant Rv3722c (10 µM) was incubated with a mycobacterial metabolite extract for 0 h or 2.5 h at 37 °C and analyzed using untargeted LC-MS. Each dot represents a feature (a chromatographic peak with a specific m/z) in the negative ionization mode; red dots represent features related to the feature with m/z 144.03 [M-H]- (n=3). **B)** Timecourse of Rv3722c-dependent formation of m/z 144.03. Same as A, but data shown for 0, 1 and 2,5 h, in presence of active Rv3722c and heat-inactivated Rv3722c control (10 min 95 °C) (mean +/- SD of three experimental replicates (n=3) representative of at least two independent experiments). **C)** MS fragmentation spectra of the Rv3722c reaction product with m/z 144.03 and in-house synthesized ketoglutaramate at a collision energy of 10. **D)** Steady-state enzyme kinetics of Rv3722c. Purified recombinant Rv3722c (0.01 µM) was incubated with 10 mM α-ketoglutarate and increasing concentrations of amino donors at 37 °C. Glu formation was measured by RapidFire mass spectrometry and used to determine initial reaction rates. Data were fitted to Michaelis-Menten kinetics using Graphpad Prism software and are represented as mean +/- SD (n=3) (Kyn: Kynurenine). **E)** Aminotransferase reaction catalyzed by Rv3722c. See also Fig S3 and S4.

Aminotransferases are the only known enzymatic source of keto-Gln^27^. We therefore tested Rv3722c for aminotransferase activity. Because aminotransferases catalyze the reversible reaction between amino acids and keto acids, we tested the ability of Rv3722c to generate ^15^N-labeled amino acids following incubation with amine-^15^N-Gln and with the same unlabeled concentrated mycobacterial metabolite extract. This approach demonstrated a clear time- and protein-dependent formation of ^15^N-Asp and ^15^N-Glu, with matching depletion of αKG (Fig S3A). This approach conversely demonstrated depletion of Asp, His, Gln and Trp upon addition of α-ketoglutarate (αKG) (Fig S3B). These data thus annotate Rv3722c, at a class level, as an aminotransferase.

### Rv3722c functions as an aspartate aminotransferase

To define the substrate specificity of Rv3722c, we next incubated Rv3722c with a panel of 26 amino acids, using αKG, pyruvate and oxaloacetate as potential amino acceptors. Oxaloacetate and αKG, but not pyruvate, served as amino acceptors, while kynurenine (Kyn), Asp and Glu served as the best amino donors (Fig S4A). In contrast, His, Cys, Asn, and Gln exhibited minimal activity (Fig S4A-B). Formal kinetic studies identified Asp as the preferred substrate and Kyn as a weaker alternative (Fig 3D). His, Cys, Asn and Gln exhibited extremely low rates of turnover consistent with kinetic side reactivity (Fig S4C). These results thus annotate Rv3722c as an aspartate aminotransferase (AspAT; Fig 3E), with a weak, but significant, side-activity towards Kyn, as observed in AspATs from other species^28, 29^.

### The sequence of Rv3722c differs from classical aspartate aminotransferases

Aminotransferases are assigned to five taxonomic classes based on the sequence of their PLP binding domain^30, 31^. AspAT’s belong to class I, which has classically been subdivided into types Ia and Ib^32^. Rv3722c, however, belongs to a recently described and structurally distinct subclass of AspATs, designated type Ic^32^, that correspond to the poorly characterized Pfam family PF12897^9^. Members of this family are absent in humans and almost exclusively present in bacteria, where they have a limited species distribution (Fig S5)^9^.

### Rv3722c has a type Ic fold

To elucidate the structural basis of the substrate recognition by Rv3722c, we solved crystal structures of the enzyme pre-incubated with Glu or Kyn at a resolution of 2.6 Å and 2.15 Å, respectively (Table S1). The overall fold of Rv3722c is similar to that of other members of the recently described type Ic subgroup of PLP-binding proteins^32^ (Fig S6). Each monomer consists of a core domain (Leu53-Asp305) that folds into a nearly perfect α/β motif comprised of a central eight-stranded (predominantly antiparallel) β-sheet surrounded by eight α-helices. Rv3722c also encodes an N-terminal auxiliary domain (Pro8-Ser52 and Gly303-Leu423) that stacks and forms an elongated segment on top of the core domain and consists of five α-helices and an antiparallel β-sheet.

### Rv3722c binds dicarboxylic acid and aromatic substrates

The structure of Rv3722c pre-incubated with Glu demonstrated clear electron density of the amino acid in the active site located at the convergence of the auxiliary and core domain (Fig S7). Glu is engaged by both polar and non-polar interactions (Fig 4A). Canonical AspATs coordinate the side chain carboxylate group of dicarboxylic substrates through a conserved active site Arg (or Lys)^33, 34^. Our structure shows that Rv3722c also forms a salt bridge with side chain carboxylate group of Glu, but using a structurally non-homologous Arg residue at position 141.

**Figure 4.**
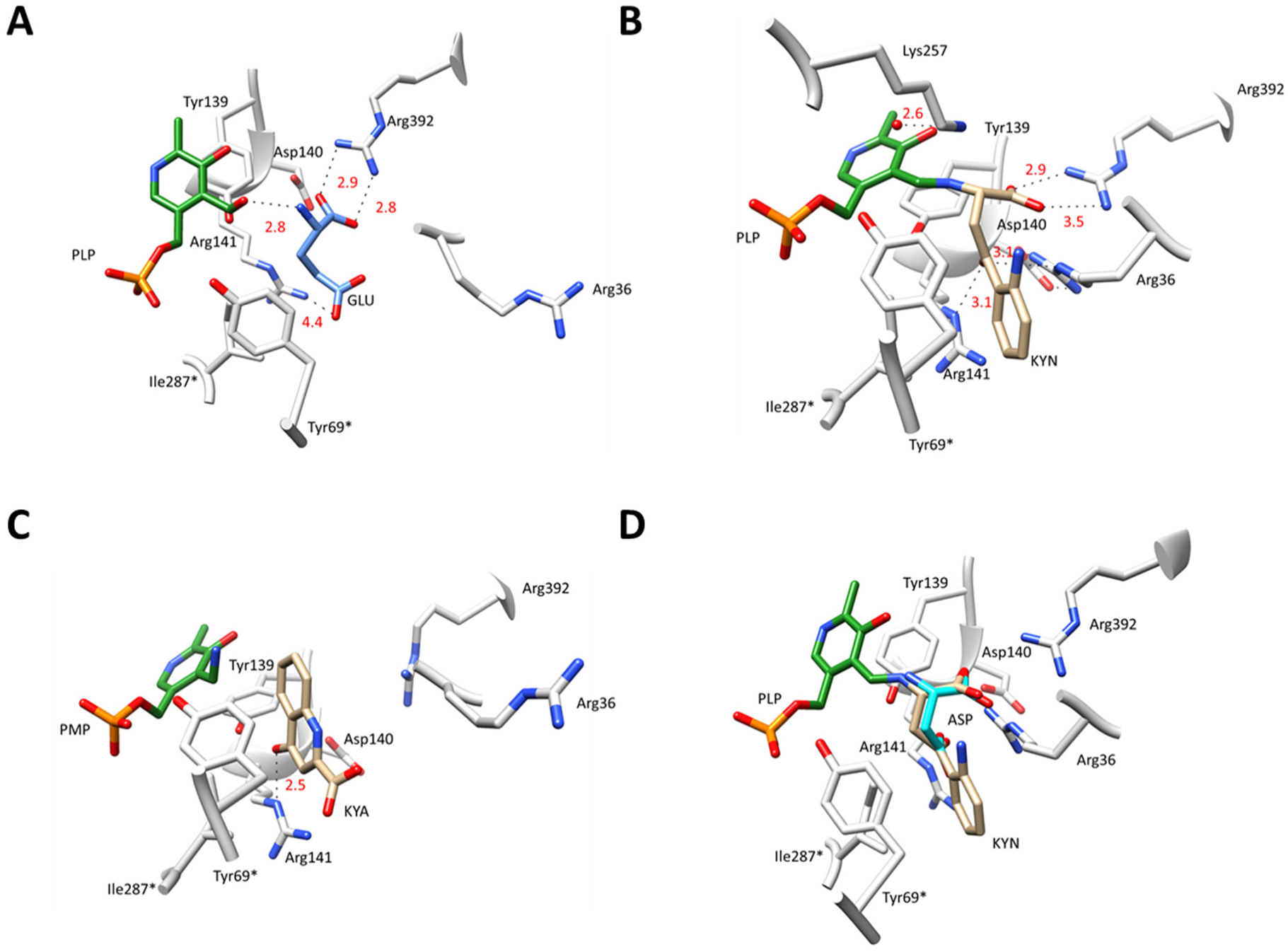
Rv3722c structure with bound ligands. **A)** Stick representation of Rv3722c in complex with glutamate (GLU; blue). **B)** Rv3722c in complex with kynurenine (KYN; tan). **C)** Rv3722c in complex with kynurenic acid (KYNA; tan). **D)** Kyn mimics Asp in the active site pocket of Rv3722c. Modelling aspartic acid (ASP; cyan) into the electron density of the aliphatic chain of KYN (tan), reveals a structural mimicry that is potentially behind the capability of Rv3722c to use KYN as an amino donor. For all figures, active site residues are in light grey and PLP in green. Ligand interactions are shown as dashed lines and distances are in angstroms. (* denotes a residue from the second monomer). See also Fig S6-S10.

In the asymmetric unit of Rv3722c pre-incubated with Kyn, we observed that the chains were bound to either the external aldimine intermediate PLP-kynurenine (PLP-Kyn), or the final keto acid product kynurenic acid (Kyna) (Fig 4B, 4C, S8A and S9). In the unbound form of Rv3722c, the positively charged guanidinium side chain of Arg36 points outward and lies above the entrance of the active site pocket (Fig S10). Upon binding Kyn, however, we observed that the side chain of this residue is positioned more inward (by ∼ 12 Å measured from the Cζ atom) and interacts with Kyn (Fig 4B, S8B and S10). Upon moving inwards, Arg36 forms a hydrogen bond salt bridge with Asp140. This ligand-induced conformationial change is reminiscent of the so called “arginine switch” observed in type Ia AspATs, although here the Arg residue moves towards the substrate rather than away^35^. In the first half reaction of transamination, a water molecule hydrolyses the ketimine intermediate releasing a keto acid of the amino donor and generating PMP^40^. In our structure, we observed a sharp peak of a water molecule in close proximity to both the active site Lys257 and the PLP-KYN intermediate (Figure S8A). We speculate that this is the same water molecule responsible for the generation of the keto acid Kyna, from the PLP-KYN intermediate.

On the other hand, we observed that the product Kyna exhibits a different pose relative to its precursor such that, when compared to Kyn, the carboxylate group of Kyna is rotated by ∼ 180° and faces the entrance of the solvent-exposed binding pocket (Fig 4C). Kyna is stabilized mainly by non-polar contacts in the binding pocket. The only major interaction stabilizing the product is a hydrogen bond between its hydroxyl group and NE group of Arg141. Interestingly, in all the Kyna-bound chains, Arg36 adopts its original “outward” conformation (Fig 4C), most likely facilitating the release of the product.

### Kyn mimics Asp in the active site pocket of Rv3722c

Modelling of Asp and superimposing it with the aliphatic chain of Kyn not only revealed that the amino groups and α-carboxylates are superimposable, but also that the carbonyl oxygen of Kyn is at a position analogous to the O6 atom of the carboxyl side chain of Asp (Fig 4D). As a result, the carbonyl oxygen is well located to hydrogen bond with Arg141 and Arg36. In addition, the binding pose observed in Kyn-bound Rv3722c allows the aromatic ring to make hydrophobic contacts with nearby residues Tyr69* and Ile287*, further stabilizing the ligand prior to transamination. Our data thus show that the aliphatic backbone of Kyn mimics the dicarboxylic amino acid Asp, and provide a possible explanation why Rv3722c – and potentially other AspAT’s – display side activity towards the structurally disparate Kyn^28, 29^.

### Rv0337c/AspC and Rv3565/AspB do not function as AspAT’s

Despite the foregoing genetic, structural, microbiological and *in vitro* biochemical evidence establishing Rv3722c as an essential AspAT, existing annotations of the *Mtb* genome include two additional AspAT’s (Rv3565/AspB and Rv0337c/AspC), one of which (Rv0337/AspC) was also predicted to be essential for *in vitro* growth^7, 8^. Unlike Rv3722c, both Rv0337c/AspC and Rv3565/AspB are annotated as class I AspATs^9, 10, 31^. We resolved this ambiguity by conducting ABMP on both enzymes. These studies demonstrated formation of keto-Gln and keto-Ile/Leu with Rv3565/AspB, and αKG with Rv0337c/AspC (Fig S11). ABMP in the presence of ^15^N-labeled amino acids and their keto acids confirmed these activities (Fig S12 and S13). These results indicate that Rv0337c/AspC encodes an alanine aminotransferase AlaT, while Rv3565/AspB encodes an alanine/valine aminotransferase AvtA with additional activity towards methionine. Both interpretations were confirmed by steady-state kinetic assays (Fig S14).

### Rv3722c is the main AspAT in Mtb

To test for other unannotated AspATs in *Mtb*, we traced the metabolic fates of ^15^N-Asp and ^15^N-Glu in Rv3722c-sufficient and -depleted *Mtb,* into ^15^N-Glu and ^15^N-Asp, respectively (Fig 5A). In both cases, we observed strictly Rv3722c-dependent transfer of the labeled amino group to the corresponding keto acid, establishing that Rv3722c is the main AspAT in *Mtb*, potentially capable of running in both directions.

**Figure 5.**
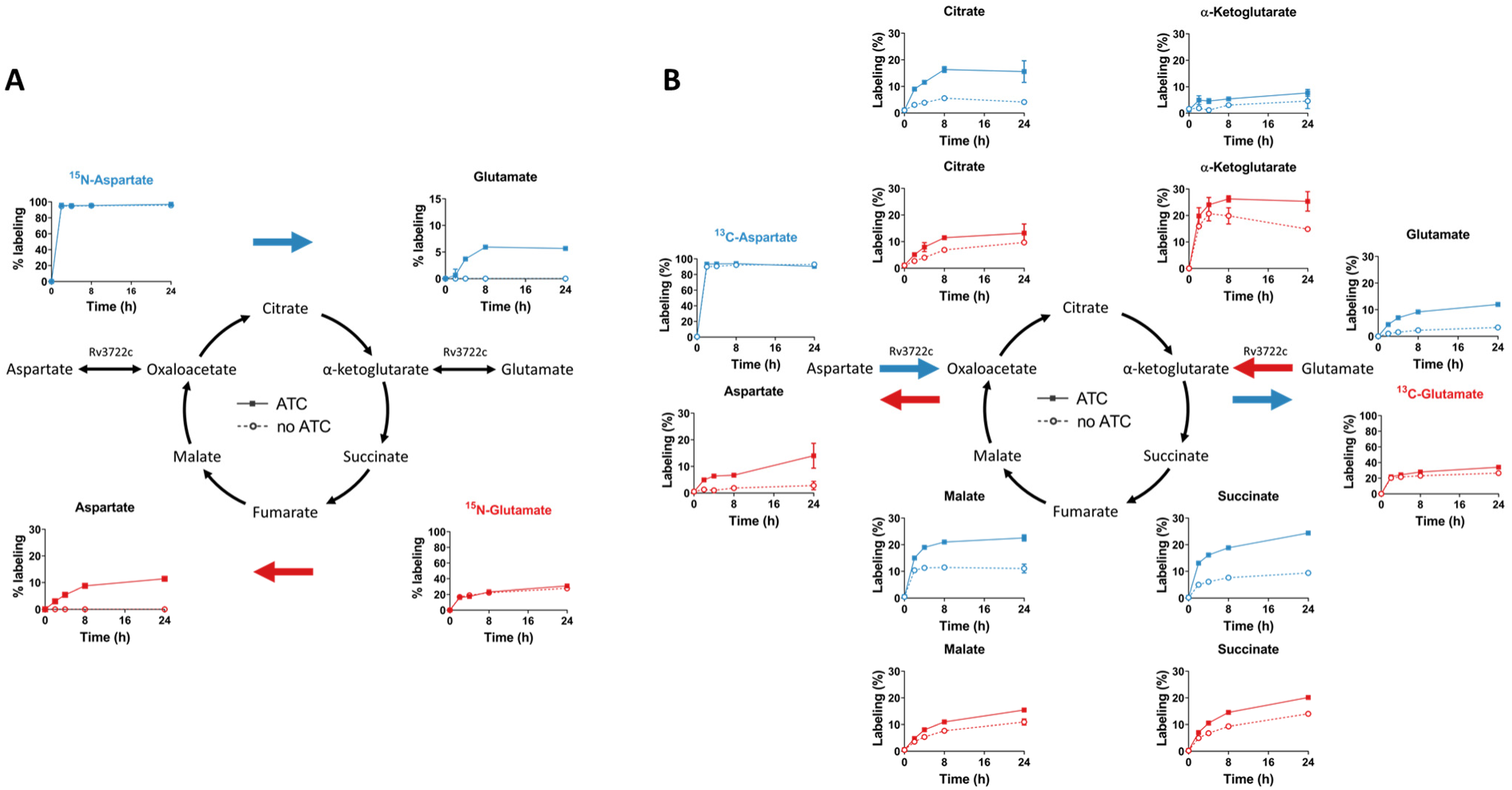
Rv3722c is the main aspartate aminotransferase in *Mtb*. **A)** ^15^N stable isotope tracing in Rv3722c-proficient and –deficient *Mtb.* Rv3722c-TetON was cultured in 7H9 supplemented with casein hydrolysate, with or without ATC, to pre-deplete Rv3722c. After a 3h adaptation to unsupplemented 7H9 with (solid line) or without ATC (dashed line), ^15^N-Asp (blue) or ^15^N-Glu (red) was added to the media at a final concentration of 3 mM. After 0, 2, 4, 8 and 24h at 37 °C, ^15^N-labeling of the indicated metabolites was determined using LC-MS. Colored arrows indicate the direction of the Rv3722c-mediated reaction. **B)** ^13^C stable isotope tracing in Rv3722c-proficient and –deficient *Mtb.* Same as (A), but using ^13^C_4_-Asp (blue) or ^13^C_5_-Glu (red). Percentages are relative to the sum of all isotopes and corrected for natural isotope abundance. Data are presented as mean +/- SD of three experimental replicates (n=3, but n=2 for ^15^N-Asp-ATC-4h, ^15^N-Glu-ATC-4h and ^15^N-Asp-noATC-2h, due to sample loss), representative of two independent experiments.

### Rv3722c balances anaplerotic and cataplerotic reactions of the TCA cycle

The conversion of Asp to oxaloacetate and Glu to αKG represent anaplerotic reactions of the TCA cycle, while their reverse reactions are cataplerotic. Rv3722c’s ability to couple one anaplerotic half-reaction of the TCA cycle with a cataplerotic counterpart suggested that its activity might help enable balanced entry and exit into and out of the oxidative and reductive arms of the TCA cycle. To test this model, we incubated *Mtb* with ^13^C-labeled Asp and Glu, and monitored for Rv3722c-dependent labeling of TCA cycle metabolites (Fig 5B). We observed that Asp could serve as carbon source for the TCA cycle in a manner primarily dependent on Rv3722c and that anaplerosis from Glu could be suppressed in the absence of Rv3722c. These results thus demonstrate that Rv3722c appears poised to mediate anaplerosis from Asp and Glu while simultaneously facilitating cataplerosis of their corresponding partner ketoacids.

### Rv3722c couples nitrogen assimilation to Asp synthesis

Given the ability of Rv3722c to couple Glu and Asp biosynthesis to one another and the widely conserved role of Glu in primary assimilation of nitrogen, we sought to define the role of Rv3722c in coupling nitrogen assimilation to Asp biosynthesis. To do so, we exposed pre-depleted Rv3722c-TetON to a nitrogen up- and downshift, in a minimal medium with ammonia as sole nitrogen source^36^. In *E coli*, nitrogen upshift causes a rapid decrease in αKG levels and slight increase in Glu levels^37^. We observed a similar, though less pronounced, effect in *Mtb*, which was not dependent on Rv3722c (Fig 6). In Rv3722c-sufficient *Mtb*, the nitrogen-dependent increase in the Glu-αKG ratio resulted in an increase in Asp synthesis, while no Asp biosynthesis was observed in Rv3722c-deficient cells (Fig 6). These results thus confirm that Asp synthesis is both strictly dependent on Rv3722c and coupled to Glu-mediated assimilation of inorganic nitrogen.

**Figure 6.**
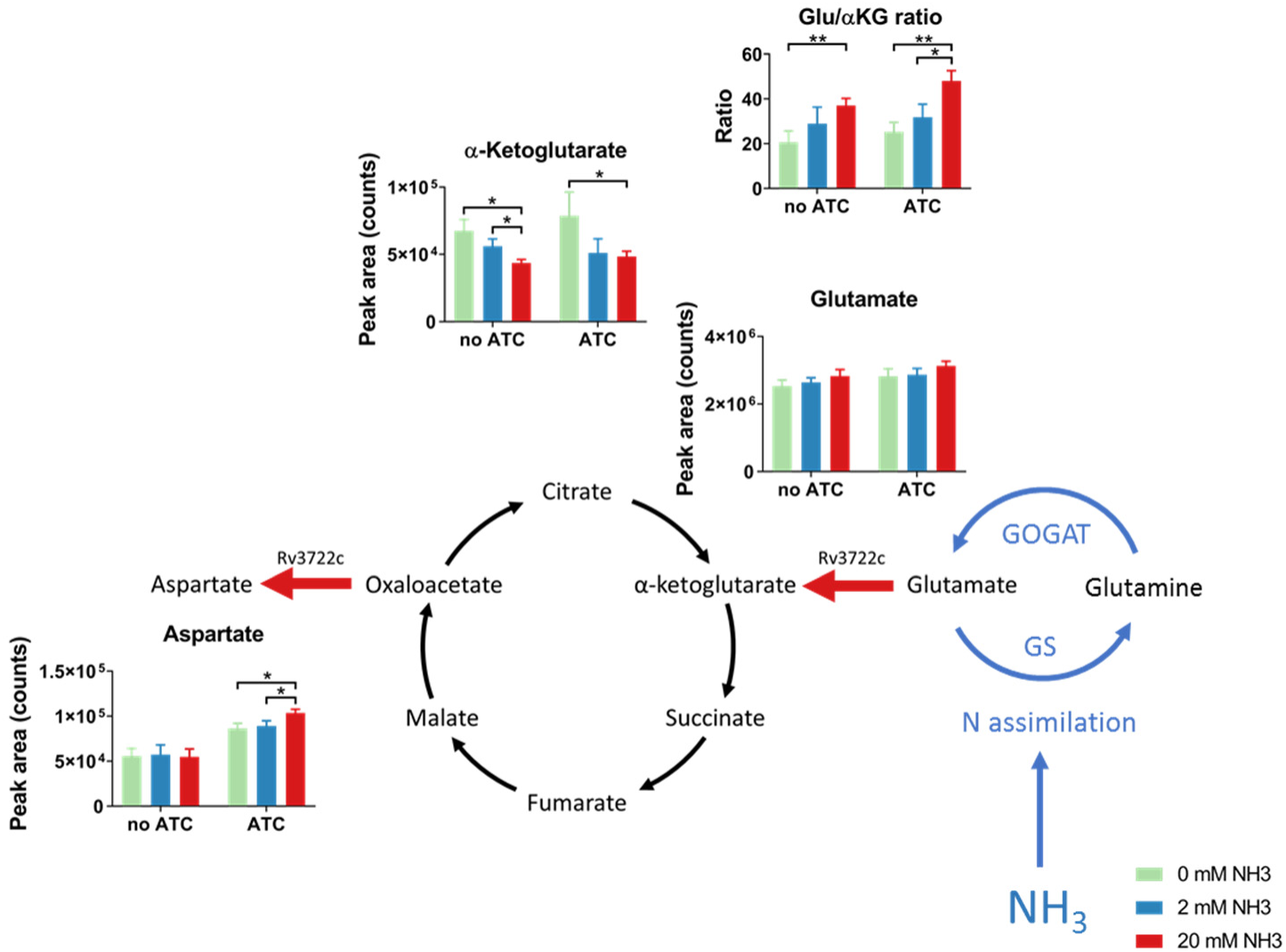
Rv3722c links nitrogen assimilation to aspartate synthesis. Rv3722c-TetON was cultured in modified minimal TSM media supplemented with 2 mM ammonium chloride and 1% casein hydrolysate, with or without ATC, to pre-deplete Rv3722c. After adaptation (24h) to the same media without casein hydrolysate and 2 mM ammonium chloride as sole nitrogen source, the media was replaced with the same media containing 0 (green), 2 (blue), or 20 mM (red) ammonium chloride. After 4h at 37 °C, the relative levels of the indicated metabolites were determined using LC-MS. Blue arrows indicate reactions involved in nitrogen assimilation (GS: glutamine synthetase; GOGAT: glutamine oxoglutarate aminotransferase), while red arrows indicate the direction of the Rv3722c-mediated reaction. Data are presented as mean +/- SD of three experimental replicates (n=3), representative of two independent experiments.

### Aspartate supplementation specifically rescues growth

Our initial growth experiments demonstrated that Rv3722c was dispensable in the presence of casein hydrolysate, a complex mixture of amino acids and small peptides (Fig 1C). We therefore sought to determine the specific amino acid(s) responsible for remedying the growth impairment of Rv3722c-deficient *Mtb* by culturing Rv3722c pre-depleted *Mtb* in 7H9 supplemented with each of 19 amino acids. Strikingly, only Asp was able to restore growth, linking Rv3722c’s *in vitro* activity to its intrabacterial metabolic activity and growth (Fig 7A).

**Figure 7.**
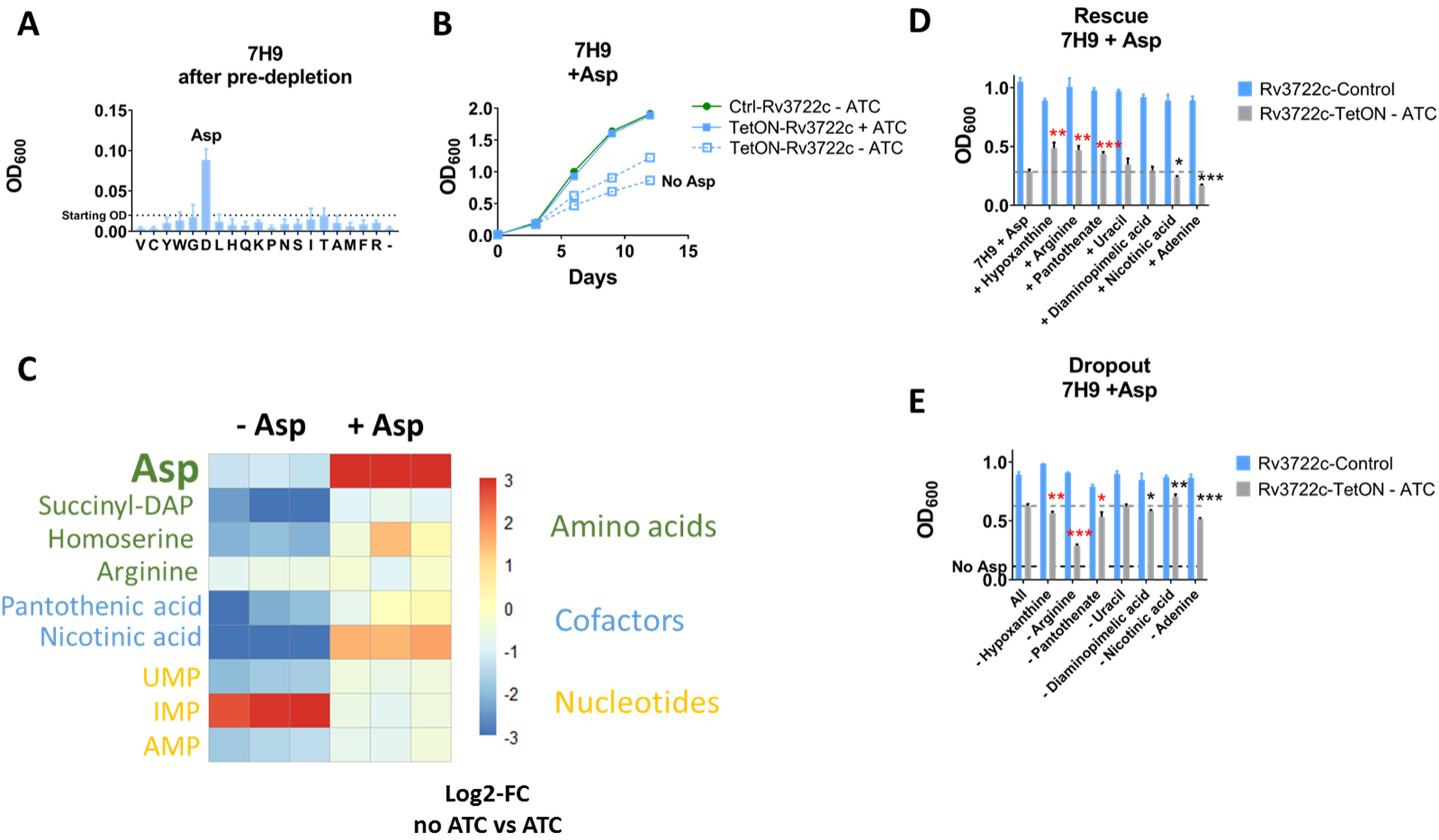
Aspartate supplementation rescues growth and metabolic defects. **A)** Rescue screen with single amino acids. Rv3722c-TetON was cultured in 7H9 with casein hydrolysate without ATC to pre-deplete Rv3722c. After predepletion, bacteria were transfered to 7H9 supplemented with 3 mM of the indicated amino acids at a final optical density at 600 nm (OD_600_) of 0.02. Bacterial growth was measured after culturing for 12 days at 37°C. Amino acids are abbreviated to their 1-letter code; -: 7H9 control. **B)** Growth curve of Rv3722c-proficient and –deficient *Mtb* in 7H9 culture media supplemented with Asp. Rv3722c-TetON and Rv3722c-control were cultured in 7H9 culture medium supplemented with 3 mM Asp, in the presence and absence of anhydrotetracycline (ATC). Data from Rv3722c-TetON cultured in the absence of ATC and Asp (Fig 1A) was plotted as a reference. **C)** Heatmap showing the levels of Asp and its essential downstream metabolites. Rv3722c-TetON was cultured in 7H9 (Fig 1A) or 7H9 supplemented with Asp (Fig 7B) with or without ATC for 12 days. Relative levels of the indicated metabolites were determined using LC-MS. Colors indicate the log_2_-fold change of the LC-MS peak areas detected without ATC versus those detected with ATC, in the absence (-) or presence (+) of 3 mM Asp. **D)** Rescue screen with essential Asp metabolites. as A, but supplementing with diaminopimelic acid, arginine, uracil (at 1 mM) or nicotinic acid, pantothenic acid, adenine or hypoxanthine (at 0.15 mM), in addition to 3 mM Asp. **E)** Dropout screen with essential Asp metabolites. Same as D, but supplementing with all but the indicated product. Bacterial growth data in A, B, D and E, and metabolite levels in F, are presented as mean +/- SD (n=3), while fold changes are presented as mean of a single ratio (n=1 per square) representative of at least two independent experiments. *: p<0.05, ** p<0.01, ***p<0.001, by a non-paired, two-tailed Student’s *t*-test (n=3), red asterisks indicate metabolites that significantly changed growth in the rescue and dropout screen. See also Fig S15.

### Aspartate is required for synthesis of growth-limiting downstream metabolites

The selective, though partial (Fig 7B), rescue by Asp suggested that Rv3722c-deficient cells suffer from a growth-limiting deficiency of Asp. To confirm this deficiency and explore its consequences on *Mtb* metabolism, we compared the metabolic profiles of Rv3722c-sufficient and -deficient bacteria, in the presence and absence of exogenous Asp. We focused on metabolites that are formed from Asp by enzymes that are essential for *Mtb*^7, 8^. Figure 7C depicts the changes related to Asp and its essential downstream products, while the full metabolite profiles collected under these and other growth conditions are shown in Fig S15. In the absence of exogenous Asp, intracellular Asp levels were reduced approximately 2-fold in the absence of Rv3722c. Levels of most essential downstream Asp-dependent products, however, were depleted to a far greater degree than their precursor, Asp (Fig 7C). Supplementation with Asp restored Asp pools to supraphysiologic levels with a similar accumulation of several downstream intermediates, such as nicotinic acid and homoserine, but corrected others, such as succinyl-DAP, less completely (Fig 7C).

To identify the specific growth-limiting, Asp-dependent metabolites among those that could not be corrected by exogenous Asp, we supplemented Asp-containing media with individual downstream metabolites. No single metabolite could completely restore wild type levels of growth. However, specific effects of pantothenate, hypoxanthine and Arg were observed in both supplementation and dropout screens (Fig 7D and E).

### Rv3722c regulates Asp-dependent distribution of assimilated nitrogen

Recognizing the ability of Asp to serve as a donor of both carbon and nitrogen^42^, we sought to determine the specific role of Rv3722c in the distribution of nitrogen via Asp. To do so, we traced the metabolic fate of the alpha amino nitrogen of Glu and Asp in Rv3722c-sufficient and -deficient *Mtb* incubated with α−^15^N-Asp or α−^15^N-Glu after 24h. As expected, we observed ^15^N labeling of known Asp-dependent metabolites following incubation with α−^15^N-Asp (Fig 8A). Incubation with α−^15^N-Glu resulted in a similar, but Rv3722c-dependent, pattern of ^15^N labeling (Fig 8A), demonstrating that Rv3722c plays a non-redundant role in the metabolic distribution of assimilated nitrogen.

**Figure 8.**
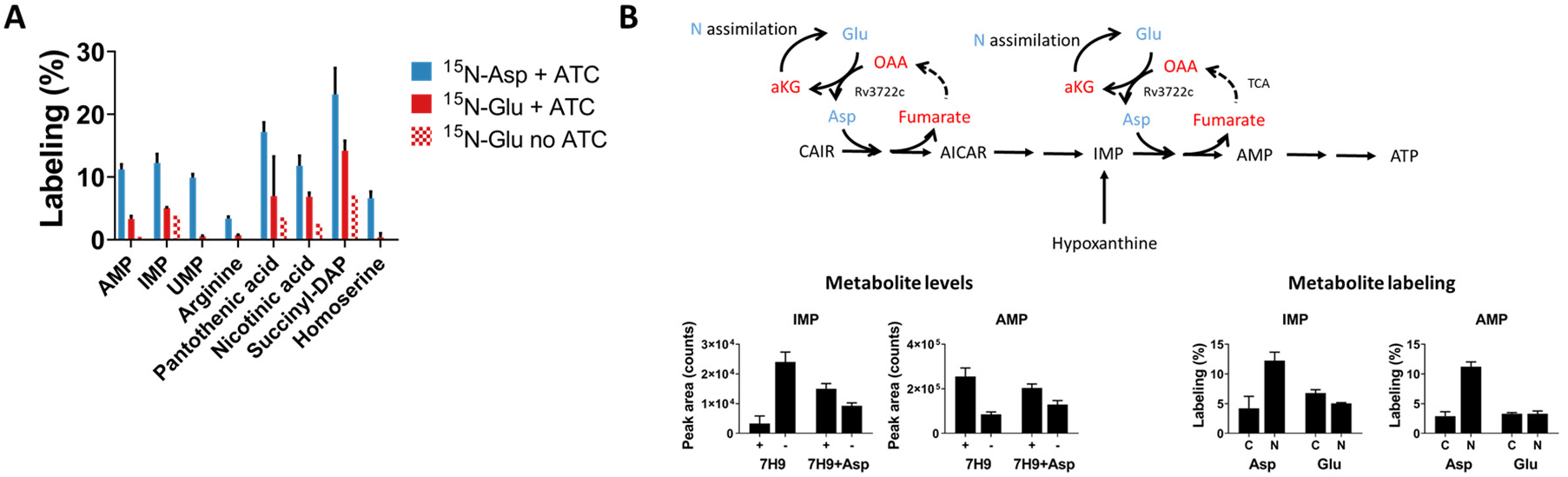
Rv3722c governs Asp-dependent nitrogen metabolism. **A)** ^15^N stable isotope tracing in Rv3722c-proficient *Mtb.* Rv3722c-TetON was cultured in 7H9 supplemented with casein hydrolysate with and without ATC. After a 3h adaptation to unsupplemented 7H9, ^15^N-Asp (blue) or ^15^N-Glu (red) was added to the media at a final concentration of 3 mM. After 24h at 37°C, ^15^N-labeling of the indicated metabolites was determined by LC-MS analysis. **B)** Rv3722c serves as a specific nitrogen conduit for *de novo* purine synthesis. Same as A, but using ^15^N and ^13^C labeled Asp and Glu. Nitrogen containing metabolites are depicted in blue, while their carbon backbones are depicted in red. Data are presented as mean +/- SD of two or three experimental replicates (n=2-3), representative of two independent replicates. OAA: Oxaloacetic acid; CAIR: Carboxyaminoimidazole ribotide; AICAR: 5-Aminoimidazole-4-carboxamide ribonucleotide.

Among Asp-dependent metabolic intermediates, we noted that levels of the purine nucleotide IMP were particularly elevated in Rv3722c-deficient *Mtb* with corresponding reductions in the levels of more downstream adenylate nucleotides, both changes being responsive to exogenous aspartate (Fig 7C and 8B). Moreover, we noted growth of Rv3722c-deficient *Mtb* supplemented with Asp could be further augmented with the addition of hypoxanthine but not adenine. Hypoxanthine is a purine salvage pathway intermediate that bypasses one of the two Asp-dependent steps in adenylate nucleotide biosynthesis (Fig 8B). Metabolic tracing similarly revealed a selective incorporation of ^15^N, but not ^13^C, into adenylate nucleotides in Rv3722c sufficient-, but not –deficient, *Mtb* (Fig 8B). These results identify a metabolically essential and specific role for Asp-derived nitrogen, unlinked to its carbon backbone, in *Mtb* growth.

## Discussion

Despite the advent of high throughput sequencing technologies, access to the biological information encoded within most genomes remains heavily dependent on sequence homology-based inference. While powerful, this approach has proven least effective for genes encoding the potentially most specific or unique biological functions of a given genome. Such approaches introduce an often unrecognized bias towards annotation of widely conserved, rather than species-specific functions^1^. Here, we applied ABMP to annotate the function of one essential and previously unannotated gene and two previously misannotated genes predicted to encode the same function. Our work highlights that the protein annotation gap is bigger than the number of unannotated proteins and includes a generally under-recognized degree of misannotation.

Our study highlights two types of misannotation, one in which two genes were erroneously annotated as AspATs on the basis of a moderate of degree of sequence homology, and another in which the identity of the actual AspAT was unrecognized. In retrospect, the misannotation of Rv0337c/AspC and Rv3565/AspB as AspATs can be explained by structural studies demonstrating that the amino acid sequence of aminotransferases is dominated by their overall fold, rather than active site-specific, architecture^38–40^. Accordingly, while the *Mtb* genome is bio-informatically predicted to encode several aminotransferases^41^, only a handful have been biochemically characterized^42–44^.

The structure of Rv3722c, in contrast, differs from canonical AspAT’s, and belongs to a recently proposed new subgroup of AspAT’s^32^. Members of this subgroup all contain an uncommon auxiliary N-terminal domain and belong to the relatively uncharacterized protein family PF12897, members of which are almost exclusively present in bacteria^9, 32^. Though the function of the auxiliary N-terminal domain is unclear, its restricted species distribution and absence in humans represent an opportunity for selective targeting of an enzyme class for which few selective inhibitors currently exist^45, 46^.

Through a combination of *in vitro* biochemistry, *in vivo* metabolomics and culture experiments, we now unambiguously resolve Rv3722c as the primary AspAT of *Mtb*, Rv0337c as an alanine transaminase and Rv3565 as an alanine-valine transaminase, further analyses of which may improve future sequence-based predictions.

Gene annotations aside, growing evidence has implicated nitrogen metabolism as an essential determinant of *Mtb*’s pathogenicity^19^. Mutants for the asparaginase AnsA, for example, exhibited impaired Asn assimilation and were attenuated in macrophages and mice^47^, while mutants for AnsP1 were unable to import Asp *in vitro* and were attenuated in mice^18^. Inhibition of glutamine synthetase, a key enzyme in nitrogen assimilation, has similarly been shown to be essential for growth in macrophages and guinea pigs^48^. Together, these studies have established nitrogen assimilation and uptake as essential determinants of *Mtb*’s pathogenicity.

Our discovery of Rv3722c as the primary AspAT of *Mtb* and its essentiality *in vitro* and *in vivo* now expands this list to include nitrogen distribution. Aminotransferases are pyridoxal phosphate (PLP)-dependent enzymes that catalyze the reversible transfer of nitrogen from an amino donor to a keto acid amino acceptor. Aminotransferases are, therefore, indispensable for the liberation and distribution of nitrogen from amino acids. The aminotransferases involved in nitrogen distribution and their essentiality in *Mtb*, however, remain largely uncharacterized^49^.

*In vitro*, aminotransferases have been shown to have somewhat overlapping substrate specificities, and *in vivo*, loss of one aminotransferase can often be overcome by artificial overexpression of another^50, 51^. In *E coli*, for example, only strains lacking both aminotransferases AspC and TyrB are Asp auxotrophs^52^. Aminotransferases have thus come to be viewed as physiologically promiscuous enzymes that catalyze highly redundant reactions^50, 51^.

Our studies indicate that *Mtb* has evolved with Rv3722c functioning as its sole AspAT in Glu-containing environments. Glu is the most prevalent metabolite in many mammalian tissues and bacteria^53–55^. Together with Gln, Glu is the primary portal of nitrogen assimilation in most bacteria^56^. Even though Asp is not a part of nitrogen assimilation, an estimated 27% of all assimilated nitrogen is distributed via Asp for the dedicated biosynthesis of several cofactors, nucleotides and amino acids^57^. Using Glu-based 7H9 culture medium, we show that Rv3722c transfers nitrogen from Glu to Asp, thus linking nitrogen assimilation to Asp-mediated nitrogen distribution. Synthesis of Asp was required for synthesis of key downstream metabolites and *in vitro* growth (Fig 1A and 5A). Rv3722c-deficient *Mtb* specifically showed depletion of these downstream metabolites, most of which could be rescued by Asp supplementation (Fig 7C). While Asp supplementation in 7H9 did not completely rescue growth (Fig 7B), the ability of more downstream Asp-dependent products to augment growth beyond that achieved with Asp alone is a likely indication of a limited flux of exogenous Asp through downstream metabolic pathways. Our results nonetheless demonstrate a non-redundant and essential role for the AspAT activity of Rv3722c.

Previous work suggested that *Mtb* has access to Asp in host macrophages^58^ and that Asp serves as an important nitrogen source *in vivo*. However, mutants deficient in the sole Asp importer Ansp1 were only partially attenuated in mice^18^. The severe attenuation of Rv3722c-deficient *Mtb* in mice suggests that access to Asp is insufficient for *Mtb* to survive during infection in mice and that the main role for Rv3722c lies in Asp biosynthesis.

We also found that Rv3722c produces Asp at a rate that is governed by the αKG-Glu ratio, which is an index for the carbon-nitrogen status. It has recently been found that *Mtb* has an extracellular Asp sensor (GlnH) that regulates the carbon-nitrogen status via GlnX, PknG and finally GarA^59, 60^. Moreover, *Mycobacterium smegmatis* strains deficient for PknG or GarA show markedly altered Asp levels^60^. Taken together, these results support a model in which Rv3722c produces Asp at a rate that is co-regulated by the presence of extracellular Asp through the αKG-Glu ratio. Moreover, our results suggest a pivotal role for Asp biosynthesis, in addition to uptake, for full virulence of *Mtb*.

Through the annotation of Rv3722c as the main AspAT in *Mtb* and the link between nitrogen assimilation and Asp biosynthesis on the one hand, and the characterization of the multifaceted essentiality of Asp on the other, we have established Asp biosynthesis as a new potential drug target in *Mtb*.

## Methods

### Chemicals

All chemicals were from Sigma, unless stated otherwise. All amino acids were in the L-form, unless stated otherwise.

### Strains

To create an *Mtb* strain with reduced levels Rv3722c, we employed a protein degradation system previously described^21^. Briefly, a FLAG epitope followed by a DAS+4 tag was recombineered into the chromosome of *Mtb* H37Rv, at the 3’-end of the target gene (H37RvMA::Rv3722c-FLAG-DAS or Rv3722c-Control). Next, the FLAG-DAS-tagged mutant was transformed with a StrepR plasmid containing *sspB* downstream of an inducible promoter (H37RvMA::Rv3722c-FLAG-DAS::sspB-pTetON-6 or Rv3722c-TetON)^20^. The *sspB-*expressing plasmid was integrated into the chromosome at the Giles phage integration site. When induced, SspB delivers DAS-tagged protein to the native protease ClpXP for degradation. Regulation was achieved by repression of the *sspB* promoter with a reverse tetracycline repressor (revTetR). RevTetR requires anhydrotetracycline (ATC), which acts as a corepressor, to shut down transcription of *sspB*. Repression of *sspB* suppresses degradation of the DAS-tagged protein. Phenotypically we thus refer to these mutants as TetON mutants. The recombineering cassette consists of 500bp flanking sequences around the stop codon of the gene, the DAS tag (inserted at the 3’-end of the target gene), a loxP site, a unique nucleotide sequence (“molecular barcode”, to enable pooled analysis of multiple strains if desired), and a hygR selectable marker. The cassette was synthesized as dsDNA (GenScript, Piscataway, NJ) in plasmid pUC57 with flanking PmeI sites. The fragment was excised from the plasmid with PmeI, and used as a double-stranded DNA recombineering substrate ^61^. Strains were confirmed with PCR and drug resistance phenotypic screening. Rv3722c-TetON was grown in the presence of 500 ng/mL ATC, which was replenished every 3-4 days. Cultures were maintained with 20 µg/mL streptomycin, but streptomycin was not added during experiments.

### Culture conditions

*Mtb* strains were cultured at 37 °C in a biosafety-level 3 facility in 10 mL standing culture flasks containing 7H9 (Middlebrook 7H9 Broth; BD) with glycerol (0.2%), sodium chloride (0.85 g/L), D-glucose (2 g/L), albumin (5 g/L; fraction V, fatty acid-free, Roche) and tyloxapol (0.04%). For specific experiments strain were grown in 7H9 with added L-amino acids (3 mM) or casamino acids (10 g/L; Hy-Case SF), or in Sauton’s (L-Asn, 4g/L; citric acid, 2 g/L; ferric ammonium citrate, 0.05 g/L; magnesium sulfate, 0.5 g/L; zinc sulfate 1 mg/L; monopotassium phosphate, 0.5 g/L; glycerol, 6%; tween 80, 0.05%; sodium hydroxide ad pH 7).

### Spot assay

Log-phase *Mtb* cultures were diluted to an OD_600_ of 0.1 in 7H9, followed by serial 10-fold dilutions. A volume of 5 µL was spotted onto square plates containing Middlebrook 7H10 agar with or without casamino acids (10 g/L; Hy-Case SF), and with or without 500 ng/mL ATC.

### Rescue with amino acids and downstream products

Rv3722c-TetON or Rv3722c-Control was precultured in 7H9 with casamino acids, without ATC, to pre-deplete Rv3722c. After washing with 7H9 without ATC, bacteria were transferred to 96-well plates containing 150 µL culture medium at a final OD_600_ of 0.02. For rescue with amino acids, medium was supplemented with 3 mM of different amino acids. For rescue with single downstream products, bacteria were transferred to media containing 3 mM Asp, and homoserine, racemic diaminopimelic acid, arginine, uracil at 1 mM, or nicotinamide, pantothenic acid, adenine and hypoxanthine at 0.15 mM. The same concentration were used for the dropout screen. Bacterial growth was monitored at 600 nm using a platereader (SpectraMax M2e, Molecular devices).

### ABMP

Activity-based metabolite profiling was performed as described^62^. In brief, a mycobacterial metabolite extract was incubated with 10 µM purified protein in 20 mM TRIS-HCl pH 7.4 or a control (heat-inactivated protein or buffer). At several time points, samples were collected into ice-cold acetonitrile:methanol:water (2:2:1; v:v:v). After centrifugation, the samples were analyzed as described under metabolomics. When described, ^15^N-labeled amino acids (10 mM; Cambridge Isotope Laboratories), or keto acids (20 mM) were added to force the reaction in one direction.

### Metabolomics

Metabolomics was performed on liquid cultures grown in media with 0.04% tyloxapol, normalized by OD_600_, after washing twice with cold PBS. Washed cells were resuspended in 1 mL ice-cold acetonitrile:methanol:water (2:2:1; v:v:v), and processed as described^62^. In brief, samples were bead-beaten (6 times for 30 s at 6500 rpm; cooled) with 0.1 mm Zirconia/silica beads (BioSpec), followed by clarification (10 min, 21,000 g, 4°C) and filter-sterilization through a 0.22 µm Spin-X filter (Sigma). Samples were stored at -80 °C until LC-MS analysis. LC-MS metabolomics was performed on non-diluted samples, using a Diamond Hydride Type C column (Cogent) on an Agilent 1200 LC system coupled to an Agilent Accurate Mass 6220 Time of Flight (TOF) spectrometer operating in the positive (amino acids) and negative (keto acids) ionization mode, as described^62, 63^.

For the detection of phosphorylated metabolites, malate, citrate and succinate, we used an ion-pairing LC-MS method. Samples were injected (5 µL) onto a ZORBAX RRHD Extend-C18 column (2.1 x 150 mm, 1.8 µm; Agilent) with ZORBAX SB-C8 (2.1 mm × 30 mm, 3.5 μm; Agilent) precolumn heated to 40 °C, and separated using a gradient of 5 mM tributylamine/5.5 mM acetate in water:methanol (97:3; v:v)(mobile phase A) and 5 mM tributylamine/5.5 mM acetate in methanol (mobile phase B) at 0.25 mL/min, as follows: 0-3.5 min: 0% B, 4-7.5 min: 30% B, 8-15 min: 35% B, 20-24 min: 99% B, 24.5-25 min: 0% B; followed by 5 minutes of re-equilibration at 0% B (Agilent 1290 Infinity LC system). Post-column, 10% dimethylsulfoximide in acetone (0.2 mL/min) was mixed with the mobile phases to increase sensitivity. Data was collected from m/z 50-1100, using an Agilent Accurate Mass 6230 Time of Flight (TOF) spectrometer with Agilent Jet Stream electrospray ionization source operating in the negative ionization mode (gas temp: 325 °C, drying gas: 8 L/min, nebulizer: 45 psig, sheath gas temp: 400 °C, sheath gas: 12 L/min, Vcap: 4000 V, Fragmentor: 125 V). Metabolites were identified based on accurate mass-retention time identifiers for masses exhibiting the expected distribution of accompanying isotopes. The abundance of extracted metabolite ion intensities was determined using Profinder 8.0 and Qualitative Analysis 7.0 (Agilent Technologies). Heatmaps were generated in R (version 3.5.2) using the pheatmap package (version 1.0.12).

### Stable isotope tracing

*Mtb* strains were cultured in 7H9 with casamino acids with or without ATC to pre-deplete Rv3722c. To accommodate large volumes, bacteria were grown in closed 1000L flasks in a shaking incubator. At the day of the experiment, the cultures were resuspended in 7H9 without tyloxapol and casamino acids at an OD_600_ of 1. After 3 hours, 3 mM ^15^N-Glu, ^15^N-Asp, ^13^C_5_-Glu and ^13^C_4_-Asp (Cambridge Isotope Laboratories) were added. After 0, 2, 4, 8 and 24 h, samples (10 mL) were collected on ice, spun down, and washed once with cold PBS. Metabolite extraction and LC-MS analysis were performed as described under metabolomics, but using Agilent Accurate Mass 6545 Quadrupole Time of Flight (Q-TOF). The percentage of labelling was determined using the batch isotopologue extraction function in Profinder software (Agilent Technologies), which corrects for natural isotope abundance. In some cases, succinyl-DAP co-eluted with and interfering signal at the M+2 m/z value. For these cases, the sample was excluded from the presented data.

### Nitrogen shift

Rv3722c-TetON was cultured in modified TSM media ^64^, containing magnesium sulfate (0.5 g/L), calcium chloride (0.5 mg/L), zinc sulfate (0.1 mg/L), copper sulfate (0.1 mg/L), ferric iron chloride (50 mg/L), monopotassium phosphate (0.5 g/L), dipotassium phosphate (1.5 g/L), sodium chloride (0.85 g/L), D-glucose (2 g/L), albumin (5 g/L; fraction V, fatty acid-free, Roche) and tyloxapol (0.04%). Bacteria were first grown in modified TSM supplemented with casamino acids (10 g/L; Hy-Case SF) and 2 mM ammonium chloride, in the presence or absence of ATC to pre-deplete Rv3722c. To accommodate large volumes, bacteria were grown in closed 1000L flasks in a shaking incubator. Next, the bacteria were transferred to the same media without casamino acids and tyloxapol, and incubated at 37 °C. After 24 h, the media was replaced with modified TSM containing 2 mM ammonium chloride (1X NH_3_), 20 mM ammonium chloride (10X NH_3_) or no ammonium chloride (0X NH_3_). Samples (10 mL) were collected, followed by centrifugation (10 min, 3000 *g*, 4°C). The pellet was resuspended in ice-cold acetonitrile:methanol:water (2:2:1; v:v:v), and processed and analyzed as described under metabolomics.

### Mouse infection

Prior to infection, Rv3722c-TetON was grown Sauton’s media with or without 500 ng/mL ATC for 10 days to pre-deplete Rv3722c. Six- to eight-week-old female BL/6 mice (Jackson Laboratory, Bar Harbor, ME) were infected in an aerosol chamber Madison chamber (University of Wisconsin). Immediately prior to infection, cells were sonicated and diluted 1:100 in PBS, and approximately 100–300 cfu administered to each mouse. Mice were fed regular chow, or chow containing doxycycline at 2000 ppm (Research Diets, New Brunswick, NJ). Lungs were harvested and plated for cfu on Middlebrook 7H10 agar or Middlebrook 7H9+1.5% Bacto Agar (Difco) containing OADC enrichment (Middlebrook) and 0.2% glycerol with ATC. Three to five mice per group were harvested at the indicated time points. To determine the number of non ATC-regulatable mutants, 2 lungs per group collected after 2, 3 and 6 weeks were plated onto plates with and without ATC. In a similar experiment, mice were infected with pre-depleted Rv3722c-TetON and wildtype, and fed chow without ATC. Lungs from wildtype mice and Rv3722c-TetON were plated on plates without ATC, or plates with and without ATC, respectively.

### Macrophage infection

Femoral mouse bone marrow cells were isolated from 7-8 week-old, male C57BL/6J mice (Jackson Laboratory) and cultured (37 °C, 5% CO2) on Petri dishes containing DMEM with 1% HEPES (Gibco), 10% FBS and 20% L cell-conditioned medium (LCM) as a source of macrophage colony-stimulating factor. On day 6, the cells were seeded in 96-well plates containing the same medium with 10% LCM at a density of 6X10^4^ cells/well. When indicated, cells were activated by adding 50 ng/mL recombinant mouse IFN gamma (Invitrogen). On day 7, the cells were infected with bacteria (cultures in 7H9 with casamino acids in the presence or absence of 500 ng/mL ATC to pre-deplete Rv3722c) in triplicate (multiplicity of infection of 1) for 4h, followed by 2 washes with PBS and the addition of fresh medium with or without 500 ng/mL ATC. On day 0, 3 and 5 post infection, the macrophages were lysed in 0.01% Triton X-100 and plated onto 7H10 plates containing 10 g/L casamino acids and 500 ng/mL ATC at serial dilutions for cfu determination. The protocol for was approved by the Institutional Animal Care and Use Committee of Weill Cornell Medicine.

### Western Blotting

Cells cultured in 7H9, 7H9 with casamino acids or Sauton’s were spun down, resuspended in 1 mL ice-cold lysis buffer (100 mM NaCl, 5% glycerol, 1 mM dithiothreitol and protease inhibitor cocktail (cOmplete, EDTA-free; Roche) in 50 mM Tris HCl, pH 8), and transferred to bead-beating tubes containing 0.1 mm Zirconia/silica beads (BioSpec). After bead-beating 3 times for 30 s at 6500 rpm in a cooled bead-beater (Precellys tissue homogenizer), samples were clarified and filter-sterilized as described under metabolomics. Samples were mixed with Laemmli sample buffer (Bio-Rad) with β-mercaptoethanol, and denatured (10 min, 95 °C) Approximately 7.5 µg of total protein (as determined by A280 on a Nanodrop; Thermo Fisher Scientific) was loaded onto 8-16% TGX gels (Bio-Rad) and run in SDS-PAGE buffer for 30 minutes at 90V followed by 45 minutes at 150 V. Proteins were transfered onto Protran 0.2 µm nitrocellulose blotting membranes (Amersham) (300 mA, 100 min, 4 °C; or 30 mA overnight, 4 °C). After washing (PBS-Tween, 30 min, room temperature) and blocking (Odyssey blocking solution; LI-COR Biosciences, 1 h, room temperature), the membranes were incubated with monoclonal mouse anti-FLAG antibody M2 (Sigma; 1:400) and rabbit anti-prcB (a gift from G. Lin and C. Nathan, 1:10,000) overnight at 4 °C. Next, the membranes were incubated with goat anti-mouse (LI-COR Biosciences, 1:20,000) and goat anti-rabbit (LI-COR Biosciences, 1:20,000) for 2 h at room temperature in Odyssey blocking buffer:PBS-Tween (1:1). After washing, proteins were detected using the Odyssey Infrared Imaging System (LI-COR Biosciences). If required, the amount of loaded protein was normalized based on the intensity of the proteasome subunit B (PrcB) loading control band in exploratory Western blots.

### Protein expression and purification

For ABMP, *Rv3722c* was cloned in a pET23(+) vector encoding a C-terminal His_6_-tag, and transformed into Bl21-AI cells (Thermo Fisher). Expression was induced at 37 °C, with 0.2 mM IPTG and 0.2% L-arabinose for 18 h. Cells were spun down, frozen at -80 °C, resuspended in 2X PBS with 10 mM imidazole, 1 mM pyridoxal phosphate, 5% glycerol, protease inhibitors (cOmplete, EDTA-free; Roche) and 1 µg/ml DNAse I, and disrupted using an Emulsiflex C5 (Avestin). After centrifugation, the lysate was loaded onto a Ni^2+^ column (HiPrep IMAC FF 16/10; GE Healthcare), followed by elution with a gradient of 10-250 mM imidazole in 2X PBS with 10 mM imidazole, 1 mM pyridoxal phosphate, 5% glycerol. Rv3722c-containing fractions were pooled and concentrated over Amicon ultra 30 KDa cutoff filter (Millipore). After buffer exchange with 50 mM Tris-HCl pH8 with 5% glycerol, the protein was loaded onto a HiPrep Q HP 16/10 and eluted with a gradient of 0-1M NaCl in 50 mM Tris-HCl pH 8 with 5% glycerol. Rv3722c-containing fractions were concentrated and the buffer was exchanged with 50 mM Tris-HCl pH 8 with 5% glycerol and 100 mM NaCl.

For structure determination, Rv3722c cloned into a modified pET28 vector encoding a TEV protease cleavage site followed by a C-terminal His_6_-tag, was chemically transformed into *E. coli* BL21 (DE3) cells. Protein production and purification was performed as outlined above, albeit, with minor modifications. Here, 0.1 mM IPTG was used to induce overnight expression at 18 °C. In addition, after Ni-NTA affinity purification, the His_6_-tag was removed using TEV protease and the protein was subjected to size exclusion chromatography (HiPrep 26/60 Sephacryl S-200; GE Healthcare) using a buffer consisting of 40 mM HEPES pH 7.4 and 150 mM NaCl. Fractions of pure Rv3722c were pooled, concentrated, flash frozen and stored at -80 °C until further use.

For ABMP, Rv0337 and Rv3565 were produced and purified using a similar approach. However, the overexpression and production of both enzymes was carried out in *E. coli* C41 (DE3) cells. Furthermore, before Rv0337 was eluted off the Ni-NTA column, an additional step, adapted and modified from^65^, was introduced to remove contaminating chaperones.

### Ketoglutaramate preparation

Ketoglutaramate was prepared and purified as described by Jaisson et al^66^. In brief, glutamine was oxidized by *Crotalus adamanteus* L-amino acid oxidase in the presence of catalase. After protein precipitation, the reaction mixture was purified over an AG 50W-X8 column (Bio-Rad), neutralized, and dried under a vacuum. The MS/MS fragmentation spectra were obtained using the LC-MS method described under metabolomics, but using an Agilent accurate mass 6545 Quadrupole-Time of Flight (Q-TOF) spectrometer operated at a collision energy of 10, 20 and 40.

### Enzyme activity assays

Substrate screens for Rv3722c were performed by incubating 1 µM protein, 1 mM amino acid, and 10 mM keto acid (αKG, oxaloacetate or pyruvate) in 150 µL 10 mM Tris-HCl pH 7.4 containing 10 µM pyridoxal phosphate at 37°C in triplicate. Samples were collected at 0, 5, 10, 30 and 60 minutes and quenched in ice-cold acetonitrile:methanol (1:1) spiked with 0.1 mM ^13^C_5_,^15^N-L-glutamic acid, ^13^C_4_,^15^N-L-aspartic acid or ^13^C_1_-L-alanine (Cambridge Isotope Laboratories) as internal standard for reactions with αKG, oxaloacetate or pyruvate, respectively. Samples were stored at -80 °C until RapidFire analysis. Rv3722c enzyme kinetics were determined by incubating purified protein (0.01 µM for L-Asp and L-Kyn, 1 µM for other substrates) with 10 mM αKG and amino acid concentrations ranging from 0.05 to 20 mM in 10 mM Tris-HCl pH 7.4 containing 10 µM pyridoxal phosphate at 37°C in triplicate. Samples were collected at 0, 0.5, 1, 2, 5, 10 and 20 minutes and quenched in ice-cold acetonitrile:methanol (1:1) spiked with 0.1 mM ^13^C_5_,^15^N-L-glutamic acid as internal standard (Cambridge Isotope Laboratories). Samples were stored at -80 °C until RapidFire analysis. Measured initial velocities were fitted to Michaelis-Menten kinetics using Graphpad Prism 7 software.

96-well plates containing enzyme activity samples were analyzed using a RapidFire high throughput MS system coupled to a 6495 triple quadrupole mass spectrometer (Agilent). Samples were loaded onto a HILIC type H1 cartridge with acetonitrile with 0.1 % formic acid, followed by a wash with the same mobile phase. Elution was performed with 20% acetonitrile in water with 0.1 % formic acid. The settings were as follows: Aspirate: 600 ms; Load/Wash: 100 ms; Elute: 5000 ms; Re-equilibrate: 4000 ms; flow: 1.25 mL/min. The mass spectrometer was set to trace the following transitions in positive ionization mode: Glutamate: m/z 148->84, CE 16; ^13^C_5_,^15^N-Glutamate: m/z 154->89, CE 16; Aspartate: m/z 134->74, CE 10; ^13^C_4_,^15^N-Aspartate: m/z 139->77, CE 10; Alanine: m/z 90 -> 44, CE 15; ^13^C-Alanine: m/z 91 -> 45, CE 15.

Rv0337c/AspC and Rv3565/AspB enzyme kinetics were determined by a coupled reaction with lactate dehydrogenase. Purified recombinant enzymes (1 µM) were incubated with 10 mM keto acid (sodium 3-methyl-2-oxobutyrate (keto-Val), (±)-3-methyl-2-oxovaleric acid sodium salt (keto-Ile), sodium 4-methyl-2-oxovalerate (keto-Leu), 2-oxoadipic acid (keto-aminoadipic acid), α-keto-γ-(methylthio)butyric acid sodium salt (keto-Met) and αKG), and L-Ala concentrations ranging from 0 to 20 mM, in 100 mM Tris-HCl pH 7.4 containing 10 µM pyridoxal phosphate, 0.5 U/mL L-lactate dehydrogenase from rabbit muscle (Roche) and 1 mM NADH (Roche) at 37°C in triplicate. The disappearance of NADH was followed spectrophotometrically at 340 nm using a SpectraMax M2e (Molecular Devices) microplate reader. Measured velocities were fitted to Michaelis-Menten kinetics using Graphpad Prism 7 software.

### Protein crystallization and Structure determination

Rv3722c (∼40 mg/mL) was co-crystalized with ligands by sitting drop vapor diffusion at 17 °C. Rv3722c pre-incubated with 5 mM L-glutamic acid at 25 °C for an hour, was added to an equal volume of reservoir solution containing 100 mM sodium acetate pH 4.5, 200 mM Li_2_SO_4_ and 50% PEG 400. Similarly, Rv3722c pre-incubated with 10 mM L-kynurenine, was mixed with a reservoir solution containing 100 mM Na_2_HPO_4_: citric acid pH 4.2, 40 % ethanol and 5 % PEG 1000. Before data collection, the crystals were cryoprotected in the mother liquor containing 25% glycerol and flash frozen in liquid nitrogen.

Diffraction datasets for Rv3722c/L-glutamic acid were collected on beamline 23 ID at Argonne National Laboratory APS synchrotron. The data were indexed, integrated and scaled using PROTEUM3 software (Version 2016.2, Bruker AXS Inc). Data were truncated in CCP4 suite^67^ and the structure was solved by molecular replacement using the unliganded structure of the same enzyme (PDB 5C6U) as a search model in MOLREP^68^.

Data for Rv3722c/L-kynurenine were also collected on beamline 23 ID at Argonne National Laboratory APS synchrotron. Data were auto processed in XDS^69^. The unmerged data were corrected for anisotropy using the STARANISO webserver (http://staraniso.globalphasing.org/cgi-bin/staraniso.cgi). The structure was solved by molecular replacement as described above.

The models were iteratively refined in PHENIX^70^ and built manually in COOT^71^. Ligand models and dictionary files were created in ELBOW BUILDER from the PHENIX^70^ suite and fitted into the density in COOT^71^. Ligand OMIT maps were calculated using Polder Maps^72^ in PHENIX. All figures were prepared in Chimera^73^. Data collection and refinement statistics are given in Table S1. The coordinates and maps of Rv3722/Glu and Rv3722/KYN have been deposited into the Protein Data Bank under accession codes 6U78 and 6U7A, respectively.

### Phylogenetic distribution

A precomputed bacterial distribution of pfam family PF12897 was downloaded from Annotree (AnnoTree v1.1.0; GTDB Bacteria Release 03-RS86; Pfam v27.0; E-value of 0.00001)^74^ and visualized in Interactive Tree of Life^75^.

## Acknowledgements

We thank Carl Nathan for critical reading of the manuscript, and the Bill and Melinda Gates Foundation TB Drug Accelerator Program (OPP1177930), NIH Tri-I TBRU (U19-AI11143), NIH NIAID Functional Genomics Program (U19-AI107774) and the Pott’s Memorial Foundation for support.

## Author contributions

RJ performed and analyzed all western blots, enzyme kinetics, macrophage infections, *in vitro* culture and metabolomics experiments; LM, RH, RJ and BS cloned, expressed and purified aminotransferases; LM, RH and JS crystalized proteins and solved their structures; SW, KG and ER performed and analyzed the mouse infection experiments; JP generated the Rv3722c mutant; RJ, KR, KG, ER, LM and JS designed the experiments and RJ, KR, LM and JS wrote the manuscript. All of the authors have read, edited, and approved the paper.

## Supplementary Information

### Results

#### Structure solution of Rv3722c in complex with glutamic acid

The X-ray diffraction data of Rv3722c in complex with Glu was initially reduced in the tetragonal crystal system and was found to belong to the P4_2_2_1_2 space group. Molecular replacement using the ligand free structure of the same enzyme (PDB ID: 5C6U) as a search model, yielded a solution comprising of a dimer in the asymmetric unit. Attempts, however, to refine the solution were unsuccessful, as the R_free_ stalled above 40%. Analysis of the data in Xtriage in Phenix^70^, revealed that the intensity statistics significantly deviated from those expected for a good to reasonable and untwinned data. In addition, the Patterson function revealed the presence of an off-origin peak at fractional coordinates (0.500 0.500 0.388), with a height approximately 45% of the origin. It has been reported that crystal pathologies such as twinning or pseudo-symmetry, can lead to wrong space group assignment because the data have an apparent high symmetry than its actual symmetry^76, 77^. Indeed, when the data was re-indexed in the lower symmetry P222 point group, the space group was found to be P2_1_2_1_2. The structure was solved by molecular replacement as described above. The resulting solution had four molecules (two dimers) in the asymmetric unit. Subsequent analysis of the data in Xtriage revealed it displayed both pseudo-merohedral twinning and pseudo translation. As a result, the twin law (-h, l, k) was used in the final round of refinement.

#### Overall structure of Rv3722c

For the structure of Rv3722c/Glu complex, residues 3 – 422 could be built into the electron density and was refined with R_work_ equal to 19.3 % and R_free_ equal to 21.6 % (Table S1). On the other hand, Rv3722c after pre-incubation with L-kynurenine crystallized in the trigonal P3_1_ space group and diffracted to 2.15 Å. There were eight molecules in the asymmetric unit. Residues 2 – 428 could be built into electron density and was refined with R_work_ equal to 17.9 % and R_free_ equal to 21.9 % (Table S1).

The overall fold of Rv3722c is similar to that of members of the recently founded Type Ic group of PLP binding proteins^32^. Each monomer consists of a core domain (Leu53 - Asp305) that folds into a nearly perfect *α/β* motif comprised of a central eight -stranded (predominantly antiparallel) *β*-sheet surrounded by eight *α*-helices. The auxiliary domain (Pro8 - Ser52 and Gly303 - Leu423), on the other hand, stacks and forms an elongated segment on top of the core domain and consists of five *α* – helices and an antiparallel *β*-sheet. A VAST search^78^ showed that the overall structure of Rv3722c was similar to the aspartate transaminase from *Corynebacterium glutamicum* (PDB 5IWQ)^32^. The root mean square difference across 418 Cα atom pairs is 1.1 Å. The two enzymes also share a high degree of sequence homology with 55% identity and 72% similarity.

#### Rv3722c complexed with substrate glutamic acid

From the structure of the Glu pre-incubated protein we observed positive electron density in the active sites of the molecules in the asymmetric unit. We observed clear electron density of Glu along that of PLP in chain B (Fig 4A and Fig S7). The rest of the chains were only bound to PLP.

Glu binds in the active site pocket located in the convergence of the auxiliary and core domain. The amino group of Glu is in close proximity to the aldehyde group of PLP at a distance of 2.8 Å. The α-carboxylate of Glu forms hydrogen bonds with (NH1; 2.90 Å) and (NH2; 2.8 Å) groups of the guanidinium side chain of Arg392. The carboxylate side chain of Asp140, whose protonation state is unclear, is within range (3.6 Å) to interact with the α-carboxylate group of Glu. AspAT’s coordinate side chain carboxylate groups of dicarboxylic substrates through a conserved active site arginine (or sometimes a lysine)^33, 79^. In Rv3722c the corresponding residue is Arg141 and it forms salt bridges with the side chain carboxylate group Glu (NH2; 4.40 Å). Hydrophobic contacts with Tyr69* and Ile287* (* denotes a residue from the second monomer) further stabilize Glu in the binding pocket.

#### Rv3722c in complex with substrate kynurenine and product kynurenic acid

We observed different ligand bound states in the molecules making the asymmetric unit of the crystals of Rv3722c pre-incubated with L-kynurenine (Kyn) (Fig S7 and S8). For accurate ligand modelling, Polder omit maps were generated^72^. The binding pocket of chain A is bound to the external aldimine intermediate PLP-kynurenine (PLP-Kyn) while the keto acid product, kynurenic acid (Kyna), was bound in chains B, C, D and G. In chains E, F and H, however, the map in the binding pocket was deemed uninterpretable and was left unmodeled.

The binding of Kyn induced only the rearrangement of the sidechain of Arg36 in the active site pocket, compared to the ligand free structure of Rv3722c (Fig S10). In the unbound form of Rv3722c, the positively charged guanidinium side chain of Arg36 points outwardly and lies above the entrance of the active site pocket. However, upon binding Kyn, the Cα of Arg36 shifts by ∼ 1 Å and the guanidinium side chain is moved inwardly by ∼ 12 Å (measured from the Cζ atom) to interact with Kyn (Figure S10). This ligand induced conformation is reminiscent of, but serves a totally opposite function than, the so called “arginine switch” observed in Type Ia AspAT^35^. An overlay of the Kyn bound with that of the ligand free structure of Rv3722c, shows that Arg36 is not in a position to sterically clash with the bulky side chain of Kyn (Figure S10). This observation suggests that the ligand-induced movement is necessary for the recognition and stabilization of the aromatic ligand. Apart from interacting with Kyn, Arg36 is locked in position by interacting with Asp140 through a bidentate salt bridge (NH1 and Oδ2; 3.6 Å) and (NH2 and Oδ1; 3 Å).

In addition to being linked to PLP, the α-carboxylate group of Kyn forms a hydrogen bond (NH1; 2.9 Å) and a salt bridge (NH2; 3.5 Å) with Arg392. The carbonyl oxygen of Kyn is hydrogen bonded to (NE; 3.1 Å) of Arg36. The carbonyl group of Kyn is also hydrogen bonded (NE; 3.1 Å) to the side chain of Arg141. The arene ring of Kyn forms hydrophobic contacts with Tyr69* and Ile287*.

In the first half reaction of transamination, the active site lysine residue abstracts a proton from C of the external aldimine yielding a quinoid intermediate36. The same Lys residue reprotonates the cofactor to generate the ketimine intermediate, which is subsequently hydrolyzed to release the keto acid of the amino donor while generating PMP. In our structure, we observed a sharp peak of a water molecule in close proximity to both the active site Lys257 residue and PLP-KYN external aldimine (Figure S8A). It is tempting to speculate that this is the water molecule responsible to hydrolyze the PLP-KYN intermediate, generating Kyna.

Interestingly, the product Kyna adopts a different pose relative to its precursor (Fig S8). In comparison to Kyn, the carboxylate group of Kyna is rotated by ∼ 180° and faces the entrance of the solvent-exposed binding pocket (Figure 4C). Unlike Kyn, the product is engaged mainly by non-polar interactions in the binding pocket. The hydroxyl group forms a hydrogen bond with (NE; 2.6 Å) of Arg141. Interestingly, in the product bound chains, Arg36 adopts its original “outward” conformation, most likely facilitating the release of the product (Figure 4C and S8). Additional hydrophobic contacts with Tyr139, Tyr69*, Ile287* stabilize Kyna in the binding pocket.

### Proposed structural basis for Kyn recognition by Rv3722c

ABMP and enzyme kinetic studies indicated that Rv3722c had significant side activity for the aromatic metabolite L-kynurenine. To our surprise, none of the aromatic amino acids were preferred by Rv3722c as amino donors. The crystal structure of Rv3722c bound to L-kynurenine provide clues why it is a suitable amino donor over the other aromatic ligands. Firstly, it appears that Arg141 is strategically positioned to demarcate the binding pocket and restrict the pose adopted by ligands in the pocket. Indeed, the binding pose adopted by Kyn could explain why it a preferred substrate. The aliphatic backbone of Kyn mimics the dicarboxylic amino acid Asp (Fig 4D). Modelling of Asp and superimposing it with the aliphatic chain of Kyn reveals that not only are the amino groups and α-carboxylates superimposable, but also importantly, the carbonyl oxygen of Kyn is at an analogous position as the O6 atom of the carboxyl side chain of Asp (Fig 4D). As a result, this allows it form hydrogen bonds with Arg141 and Arg36. In addition, this binding pose allows aromatic ring to hydrophobic contacts with Tyr69* and Ile287*– further stabilizing the ligand prior to transamination. The presence of the carbonyl group in Kyn and the lack thereof in aromatic amino acid substrates such as Trp, Phe and Tyr, appears to be a selectivity factor for binding in the active site pocket as well as the capacity to serve as an amino donor. Indeed, we also observed that 3-hydroxykynurenine, which is structurally similar to Kyn, could be used as an amino donor by Rv3722c (data not shown) – further supporting the importance of the carbonyl group in the recognition and utilization of these aromatic ligands as substrates. Taken together, these data provide insights as to why, in addition to its cognate dicarboxylic acid substrates, Rv3722c is able to bind in the same pocket and utilize as a substrate, the structurally disparate aromatic ligand Kyn. Lastly, the high degree of residue conservation in the active sites of Type Ic AspATs, suggests that side activity towards Kyn could be a hallmark of these enzymes. However, the physiologic relevance of this side activity, especially in the context of *Mycobacterium tuberculosis*, remains unclear and worth exploring.

**Table S1.**
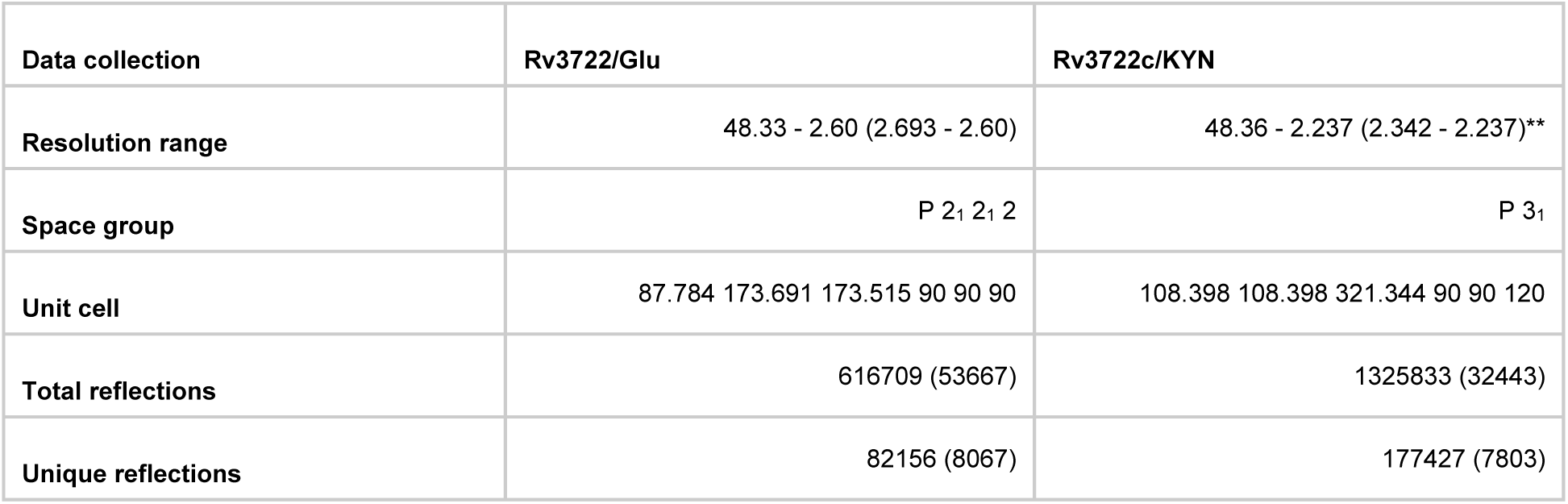
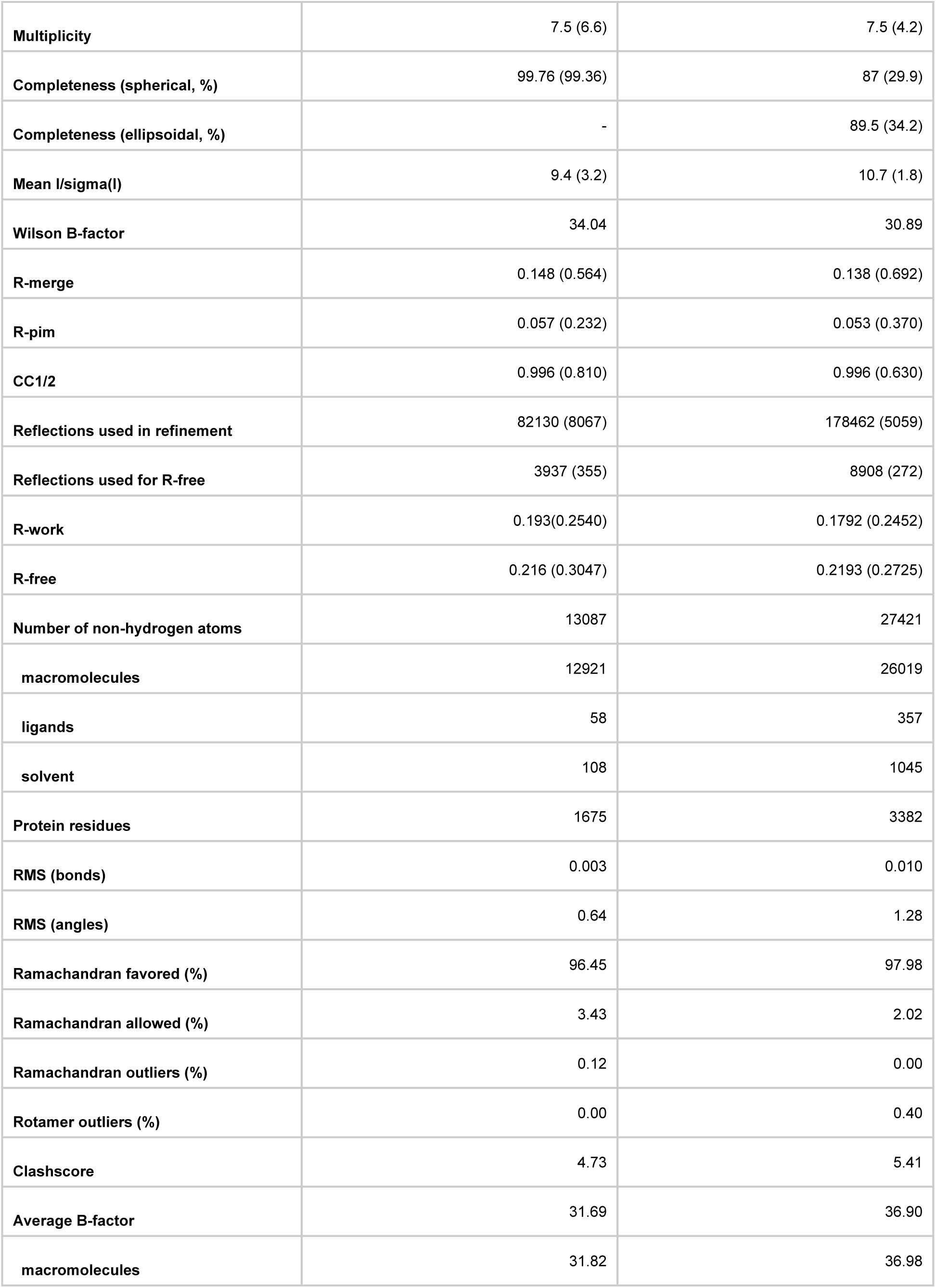
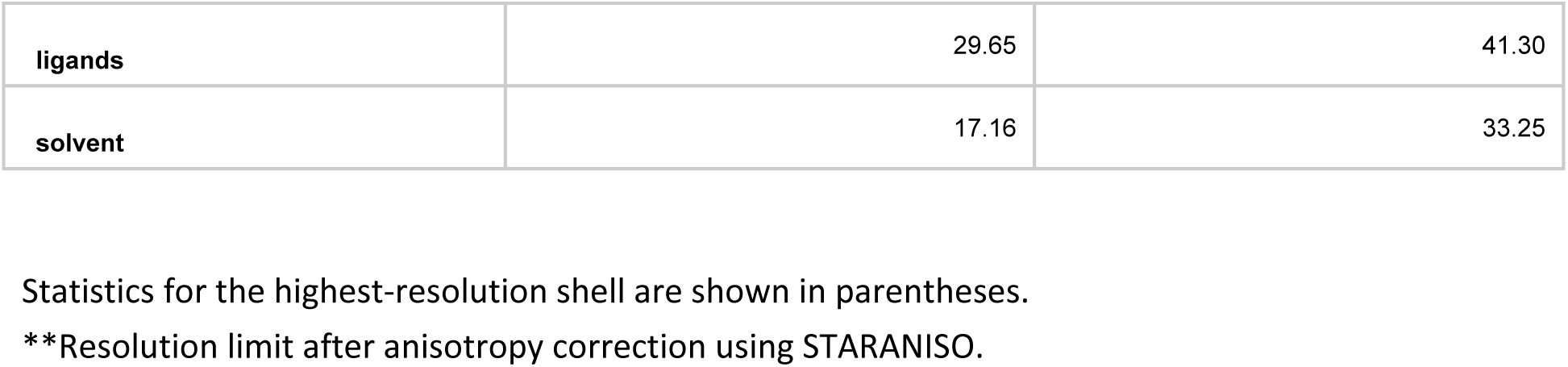
Data collection and refinement statistics.

| Data collection | Rv3722/Glu | Rv3722c/KYN |
| --- | --- | --- |
| Resolution range | 48.33 - 2.60 (2.693 - 2.60) | 48.36 - 2.237 (2.342 - 2.237)** |
| Space group | P 2<sub>1</sub> 2<sub>1</sub> 2 | P 3<sub>1</sub> |
| Unit cell | 87.784 173.691 173.515 90 90 90 | 108.398 108.398 321.344 90 90 120 |
| Total reflections | 616709 (53667) | 1325833 (32443) |
| Unique reflections | 82156 (8067) | 177427 (7803) |
| **Multiplicity** | 7.5 (6.6) | 7.5 (4.2) |
| **Completeness (spherical, %)** | 99.76 (99.36) | 87 (29.9) |
| **Completeness (ellipsoidal, %)** | - | 89.5 (34.2) |
| **Mean I/sigma(I)** | 9.4 (3.2) | 10.7 (1.8) |
| **Wilson B-factor** | 34.04 | 30.89 |
| **R-merge** | 0.148 (0.564) | 0.138 (0.692) |
| **R-pim** | 0.057 (0.232) | 0.053 (0.370) |
| **CC1/2** | 0.996 (0.810) | 0.996 (0.630) |
| **Reflections used in refinement** | 82130 (8067) | 178462 (5059) |
| **Reflections used for R-free** | 3937 (355) | 8908 (272) |
| **R-work** | 0.193(0.2540) | 0.1792 (0.2452) |
| **R-free** | 0.216 (0.3047) | 0.2193 (0.2725) |
| **Number of non-hydrogen atoms** | 13087 | 27421 |
| **macromolecules** | 12921 | 26019 |
| **ligands** | 58 | 357 |
| **solvent** | 108 | 1045 |
| **Protein residues** | 1675 | 3382 |
| **RMS (bonds)** | 0.003 | 0.010 |
| **RMS (angles)** | 0.64 | 1.28 |
| **Ramachandran favored (%)** | 96.45 | 97.98 |
| **Ramachandran allowed (%)** | 3.43 | 2.02 |
| **Ramachandran outliers (%)** | 0.12 | 0.00 |
| **Rotamer outliers (%)** | 0.00 | 0.40 |
| **Clashscore** | 4.73 | 5.41 |
| **Average B-factor** | 31.69 | 36.90 |
| **macromolecules** | 31.82 | 36.98 |
| ligands | 29.65 | 41.30 |
| solvent | 17.16 | 33.25 |

Statistics for the highest-resolution shell are shown in parentheses. \*\*Resolution limit after anisotropy correction using STARANISO.

### Supplemental Figures

**Figure S1.**
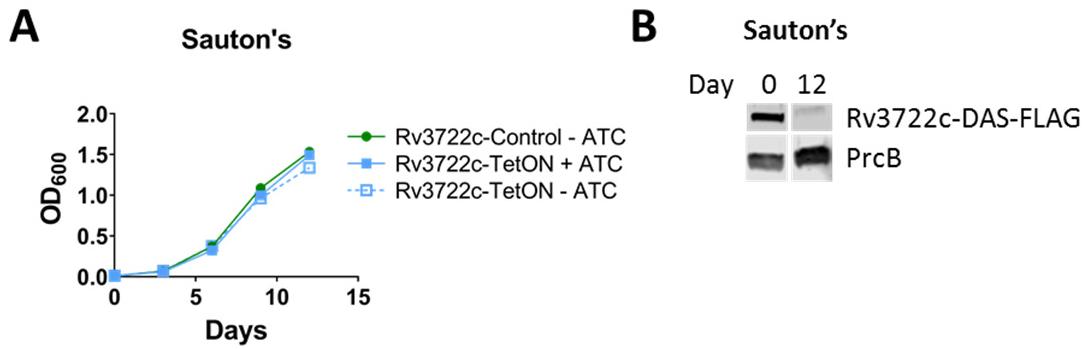
Rv3722c is dispensible in defined Sauton’s minimal media. **A)** Growth curve of Rv3722c-proficient and –deficient *Mtb* in defined Sauton’ s minimal media. Rv3722c-TetOn and Rv3722c-control cultured in Middlebrook 7H9 culture media with 500 ng/mL anhydrotetracycline (ATC) were used to inoculate Sauton’s minimal media with and without 500 ng/mL ATC. Bacterial growth was monitored for 12 days, by optical density at 600 nm. Data are represented as mean -/+ SD of three experimental replicates (n=3), representative of at least two independent experiments. **B)** Western blot showing depletion of Rv3722c after 12 days of culturing in Sauton’s without ATC (A). Protein lysates were analyzed by Western blotting, using an α-FLAG antibody. The proteasome subunit β (PrcB) was used as loading control.

**Figure S2.**
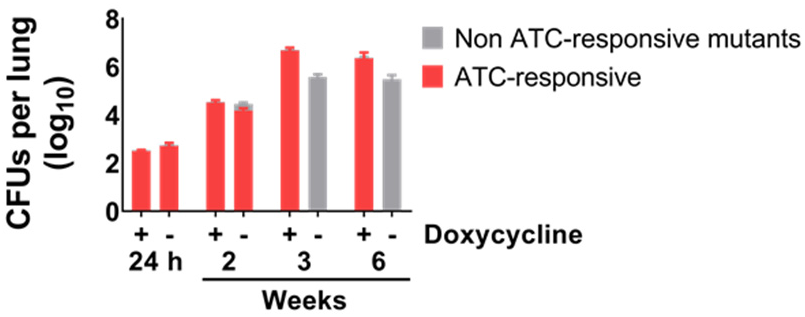
Rv3722c is required for virulence in mice. Mice were infected with aerosolized Rv3722c-TetON pre-cultured in Sauton’s with and without ATC to generate Rv3722c-proficient and -deficient *Mtb*, respectively. After infection, mice were fed chow with (Rv3722c-proficient *Mtb*) or without (Rv3722-deficient *Mtb*) doxycycline. The number of CFU’s were assessed by plating serial dilutions on 7H10 solid media with ATC. The number of non ATC-responsive mutants were determined by plating on 7H10 solid media with and without ATC (n=2). Data are represented as mean +/- SD (n=3-5). Within 3 weeks after aerosol infection of mice, Rv3722-deficient *Mtb* were almost completely replaced by escape mutants that were no longer ATC-responsive.

**Figure S3.**
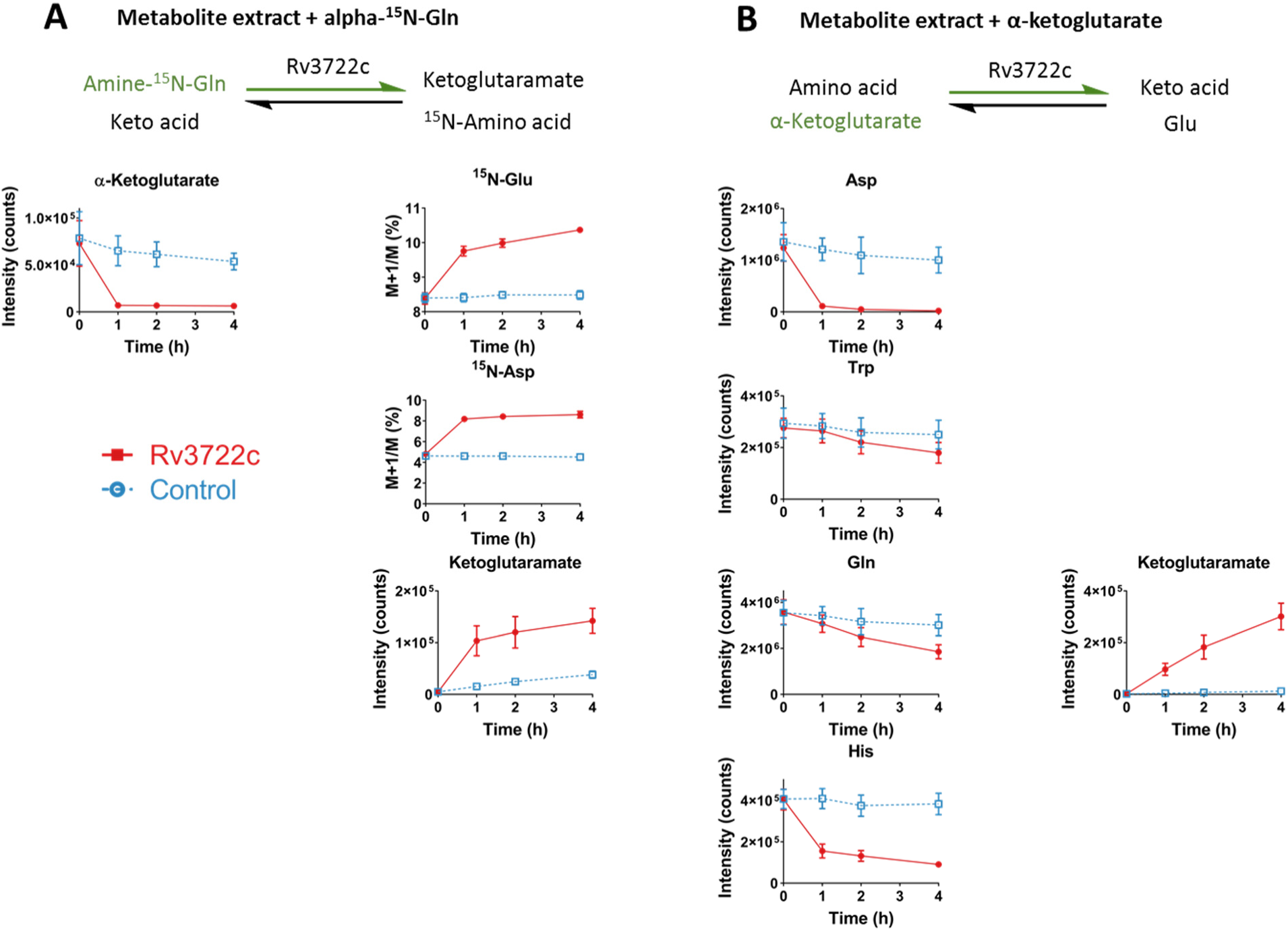
Rv3722c functions as an aminotransferase. **A)** Activity-based metabolite profiling (ABMP) with Rv3722c in the presence of α-^15^N-Gln. Purified recombinant Rv3722c (10 µM; red line) or a heat-inactivated control (10 min 95 °C; blue line), was incubated with a mycobacterial metabolite extract supplemented with 10 mM α-^15^N-Gln for 0, 1, 2 and 4 h at 37 °C, and analyzed using untargeted LC-MS. **B)** Activity-based metabolite profiling (ABMP) with Rv3722c in the presence of α-ketoglutarate. Same as A, but using a mycobacterial metabolite extract supplemented with 20 mM α-ketoglutarate. Colored arrows indicate the forced direction of the Rv3722c-mediated reaction. Relative metabolite levels are represented as intensity, while ^15^N-labeling is presented as the ratio M+1/M, which was not corrected for naturally occuring isotopes. Data are presented as mean +/- SD (n=3).

**Figure S4.**
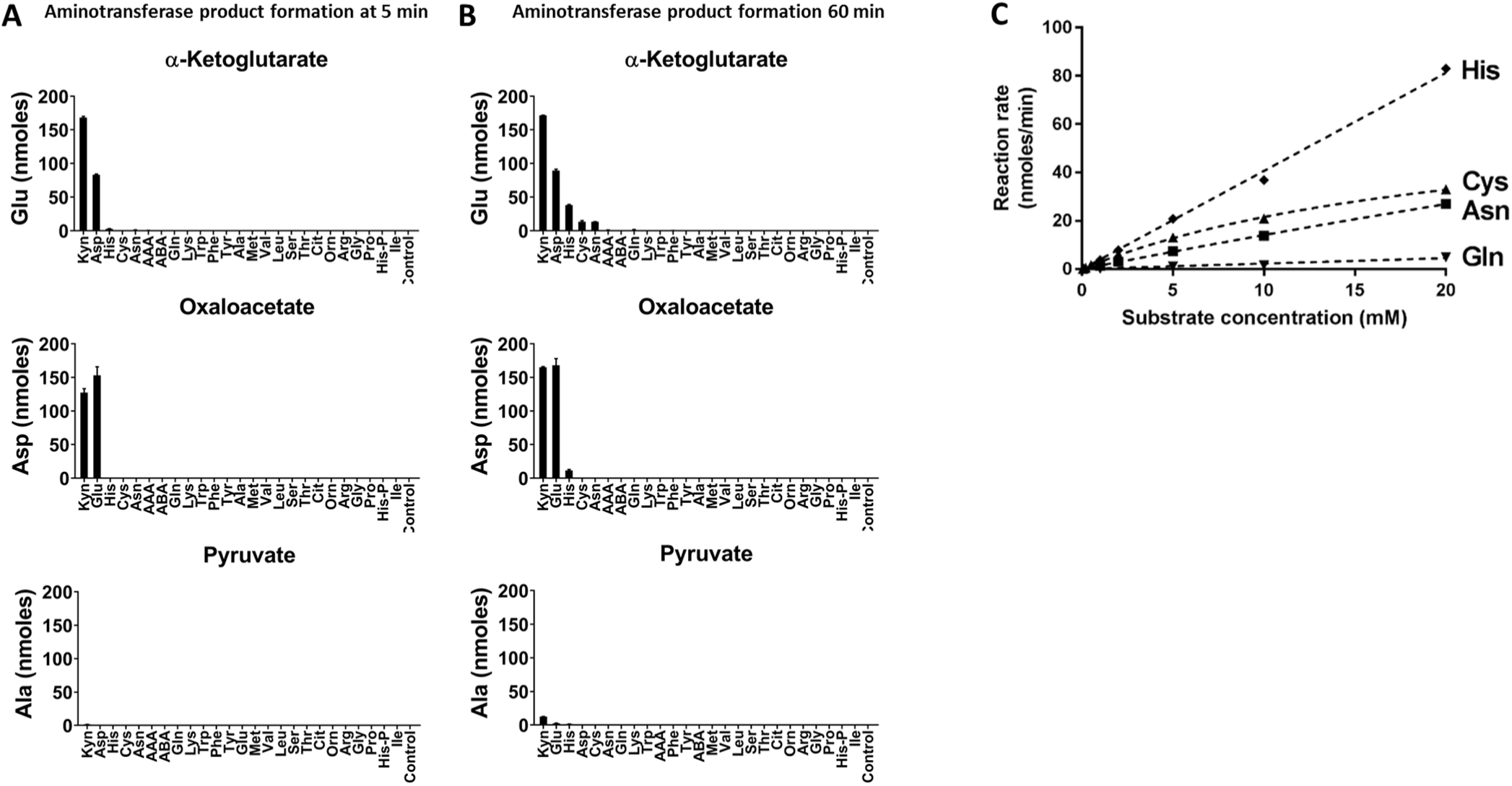
Rv3722c substrate screen. **A)** Rv3722c product formation after 5 min incubation. Purified recombinant Rv3722c (1 µM) was incubated with a panel of amino acids (1 mM) and 3 common keto acids (α-ketoglutarate, oxaloacetate or pyruvate; 10 mM). After 5 minutes at 37°C, product formation (Glu, Asp or Ala for α-ketoglutarate oxaloacetate or pyruvate, respectively) was measured by RapidFire mass spectrometry. **B)** Rv3722c product formation after 60 min incubation. Same as A, but after incubating for 60 minutes. Data are presented as mean +/- SD (n=3). AAA: aminoadipic acid; ABA: 2-aminobutyric acid; Cit: citrulline; Orn: ornithine; His-P: histidinol phosphate. **C)** Steady-state enzyme kinetics of Rv3722c for His, Cys, Asn and Gln. Purified recombinant Rv3722c (1 µM) was incubated with 10 mM α-ketoglutarate and increasing concentrations of amino donors at 37 °C. Glu formation was measured by RapidFire mass spectrometry and used to determine initial reaction rates. Data were fitted to Michaelis-Menten kinetics using Graphpad Prism software and are represented as mean +/- SD (n=3).

**Figure S5.**
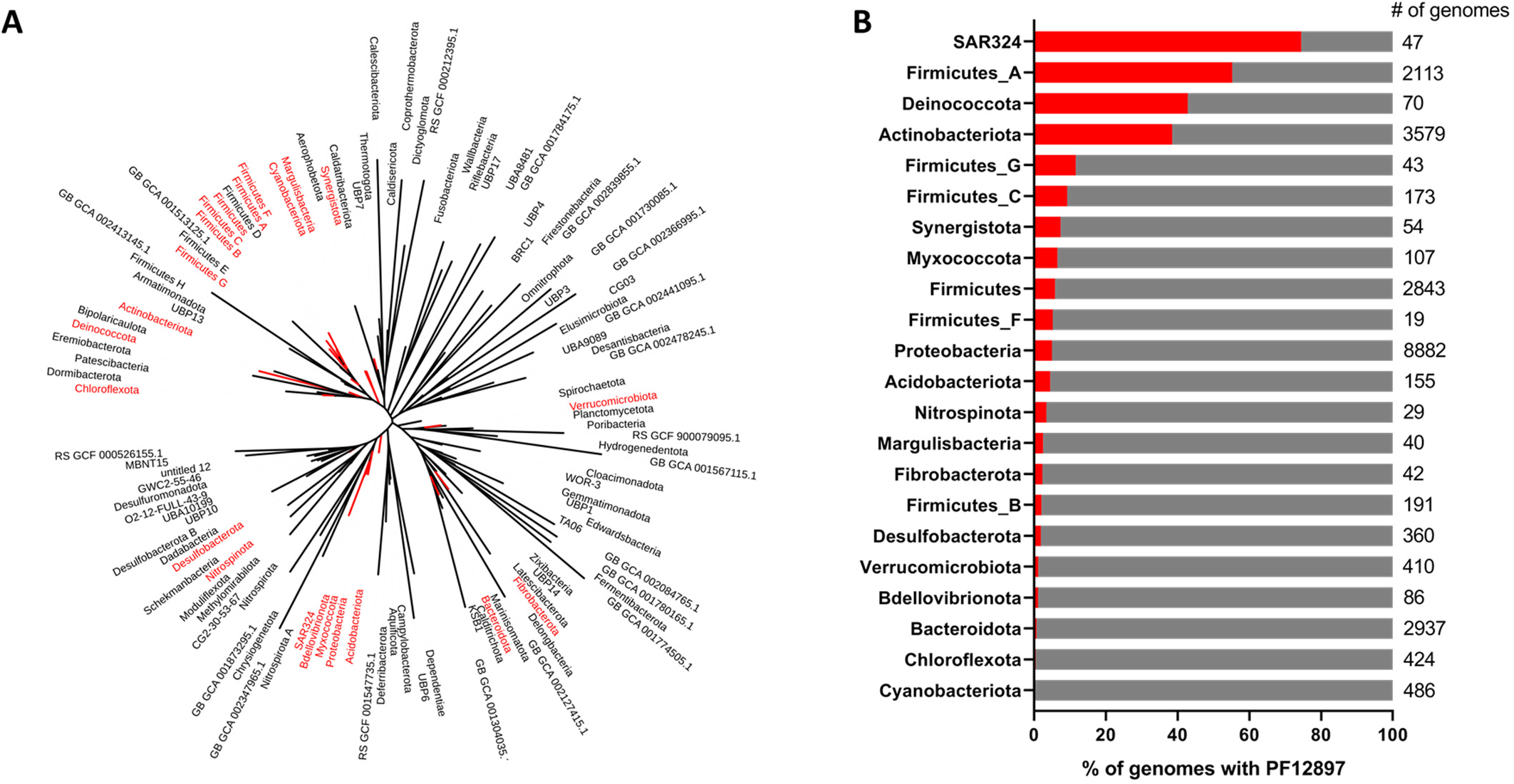
PF12897 family members are present in selective bacterial phyla. **A)** Phylogenetic tree showing the distribution of PF12897 across bacterial phyla. Phyla highlighted in red contain at least 1 genome containing a PF12897 family member. **B)** PF12897 frequency and size of phyla containing PF12897 family members.

**Figure S6.**
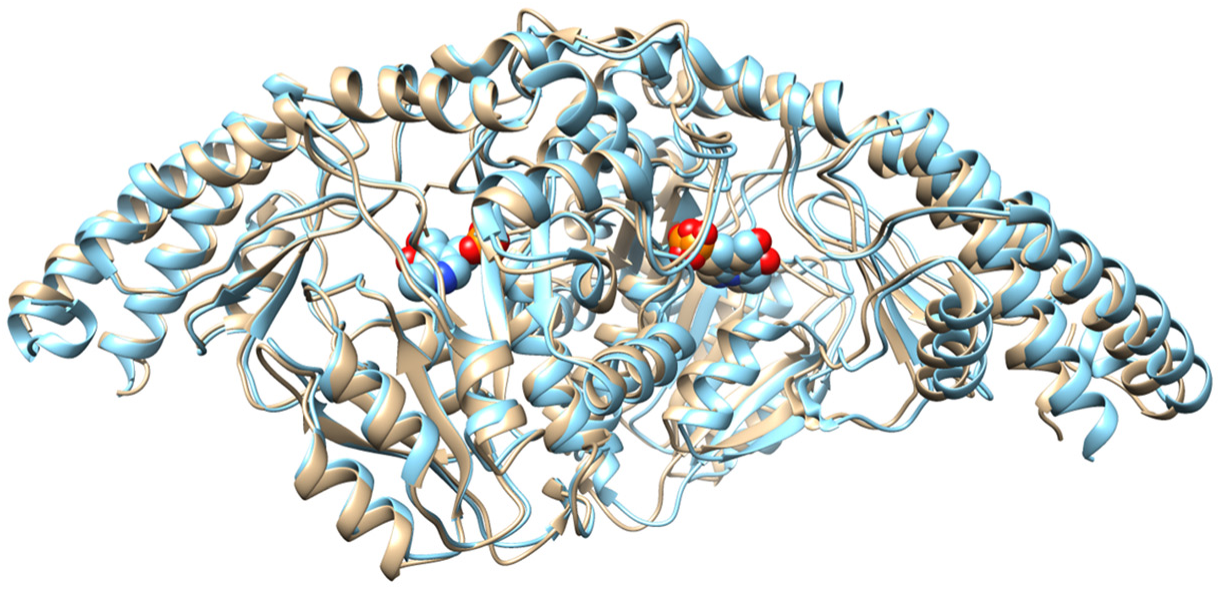
Rv3722c is a type Ic PLP-binding protein. Ribbon representation of the superposition of Rv3722c (tan) with the type Ic AspAT from *Corynebacterium glutamicum* (5IWQ; cyan). The cofactor PLP is shown as spheres in the active site pocket.

**Figure S7.**
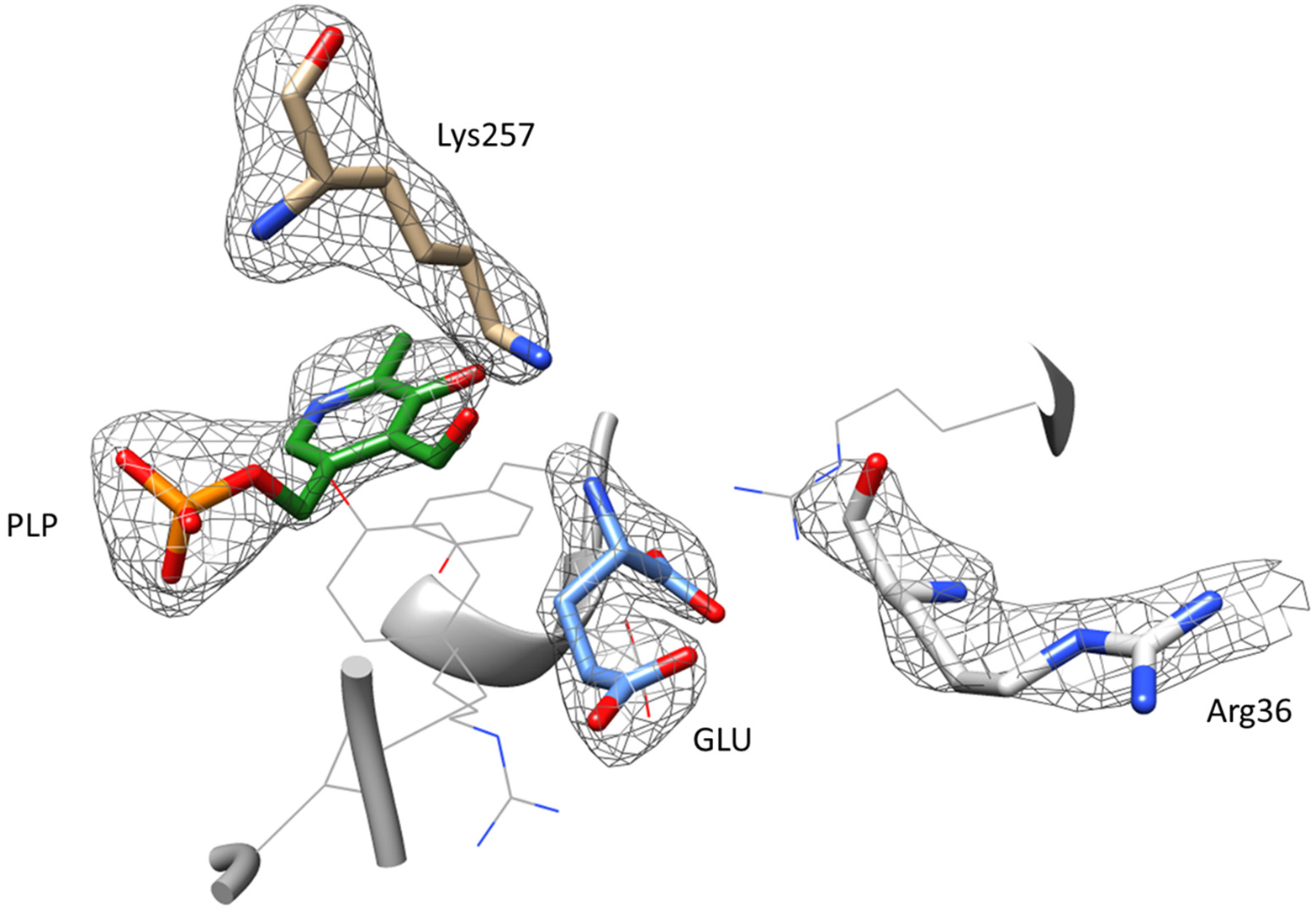
Polder omit map of glutamate in the active site of Rv3722c. The catalytic Lys257 residue is shown to highlight its position relative to pyridoxal-phosphate (PLP). Arg36 does not undergo any conformational change upon binding of GLU. The map is contoured at 3σ. Other coordinating residues are shown as wire for clarity.

**Figure S8.**
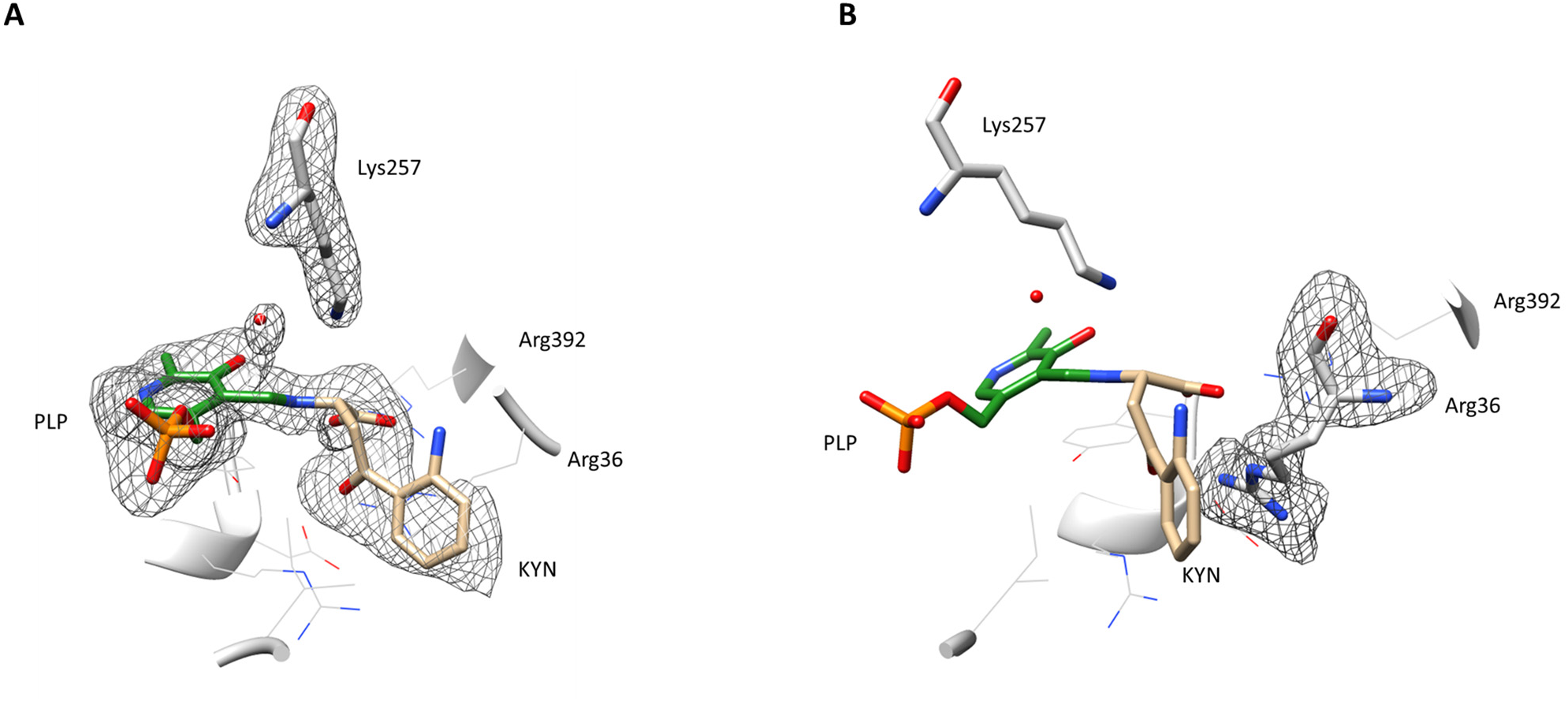
Polder omit map of the PLP-Kynurenine intermediate. **A)** The catalytic Lys257 residue is shown to highlight its clear detachment from pyridoxal-phosphate (PLP). A water molecule is found to be in close proximity to both the active site Ly257 and the PLP-KYN intermediate. **B)** Upon binding KYN, Arg36 undergoes a conformational change. The maps are contoured at 4 σ. Other coordinating residues are shown as wires for clarity. Some atoms of the arene ring of KYN did not have electron densityy.

**Figure S9.**
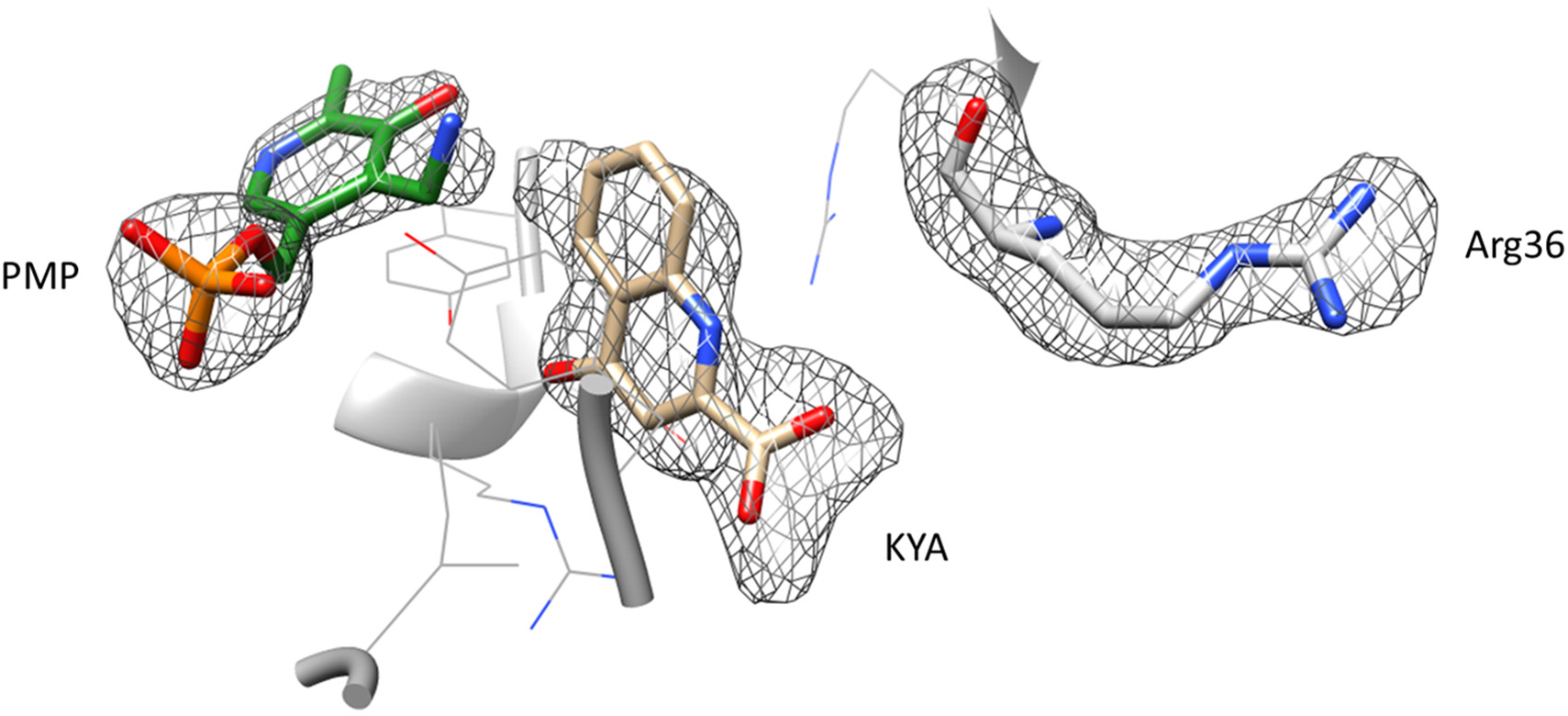
Polder omit map of PMP and kynurenic acid in the active site of Rv3722c. The product kynurenic acid (KYA) adopts a different pose relative to kynurenine. Arg36 adopts its outward conformation. The map is contoured at 4 σ. Other active site residues have been as wires for clarity. PMP: pyridoxamine phosphate.

**Figure S10.**
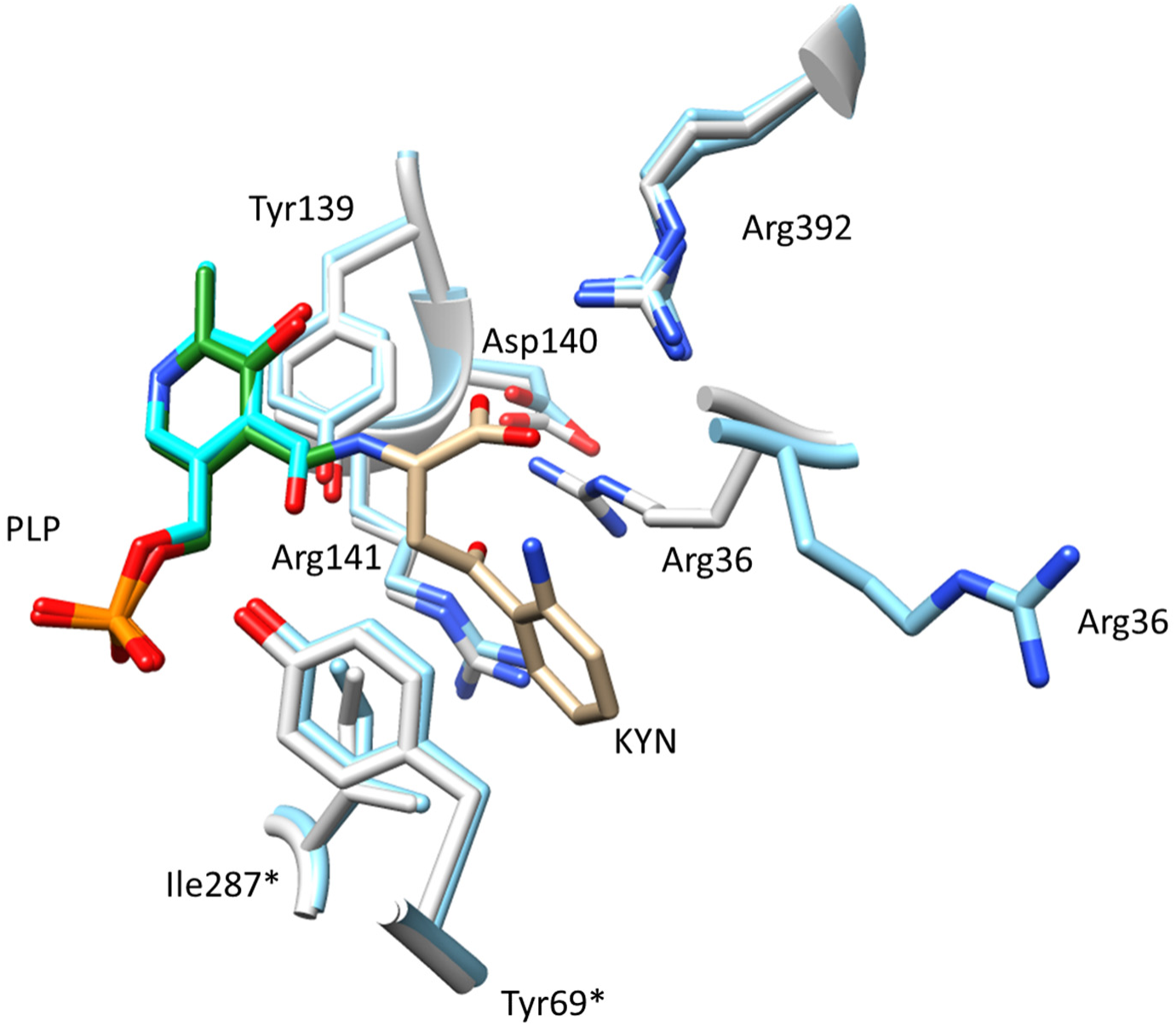
Binding of kynurenine induces a conformational change. Stick and ribbon representation of the superposition of the active sites of ligand free Rv3722c (5C6U; cyan) and PLP-KYN intermediate bound (grey) structures. The binding of kynurenine induces a conformational change of Arg36, resulting in the residue to move inwards to with the ligand.

**Figure S11.**
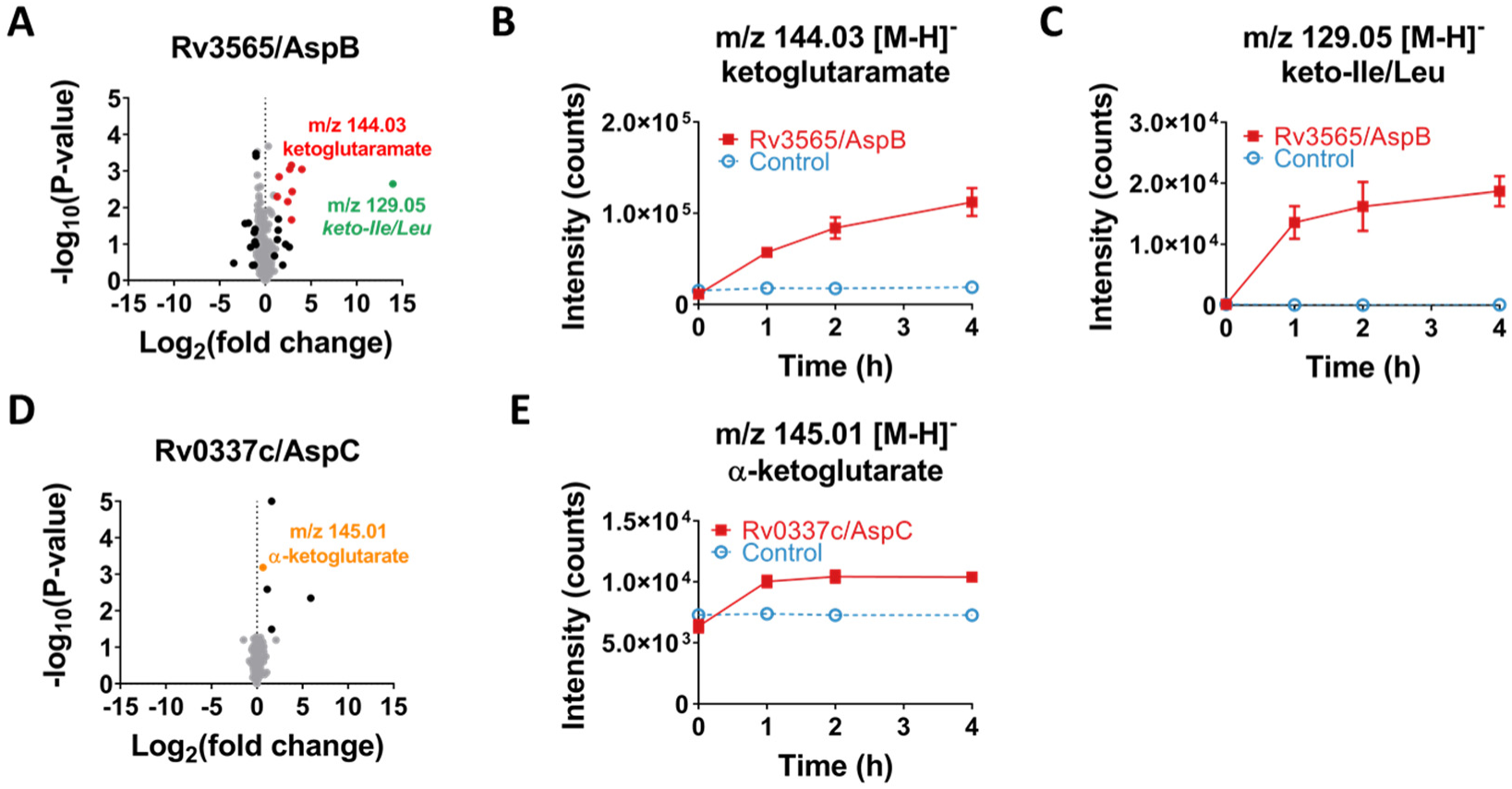
Activity-based metabolite profiling on Rv3565/AspB and Rv0337c/AspC. **A)** Volcano-plot of activity-based metabolite profiling with purified recombinant Rv3565/AspB. Purified recombinant Rv3565 (10 µM) was incubated with a mycobacterial metabolite extract for 0 h or 4 h at 37 °C, and analyzed using untargeted LC-MS. Each dot represent a feature (a chromatographic peak with a specific m/z) in the negative ionization mode; red dots represent features related (fragments, adducts, dimers and isotopes) to ketoglutaramate (m/z 144.03 [M-H]^-^), while black dots represent uncharacterized features with a fold change greater than 2 and p-value below 0.05 (n=3). **B)** Time- and Rv3565-dependent formation of ketoglutaramate. **C)** Time- and Rv3565-dependent formation of keto-Leu/Ile (m/z 129.05 [M-H]^-^; keto-Ile/Ile could not be distinguished based on mass or retentiont time). **D)** Volcano-plot of activity-based metabolite profiling with purified recombinant Rv0337c/AspC. Same as A, but using Rv0337c/AspC incubated for 0 h or 1h at 37 °C. The orange dot represents α-ketoglutarate (m/z 145.01 [M-H]^-^) while black dots represent uncharacterized features with a fold change greater than 2 and p-value below 0.05 (n=3). **E)** Time- and Rv0337c-dependent formation of α-ketoglutarate. Data in panel B, C and E are presented as mean +/- SD (n=3).

**Figure S12.**
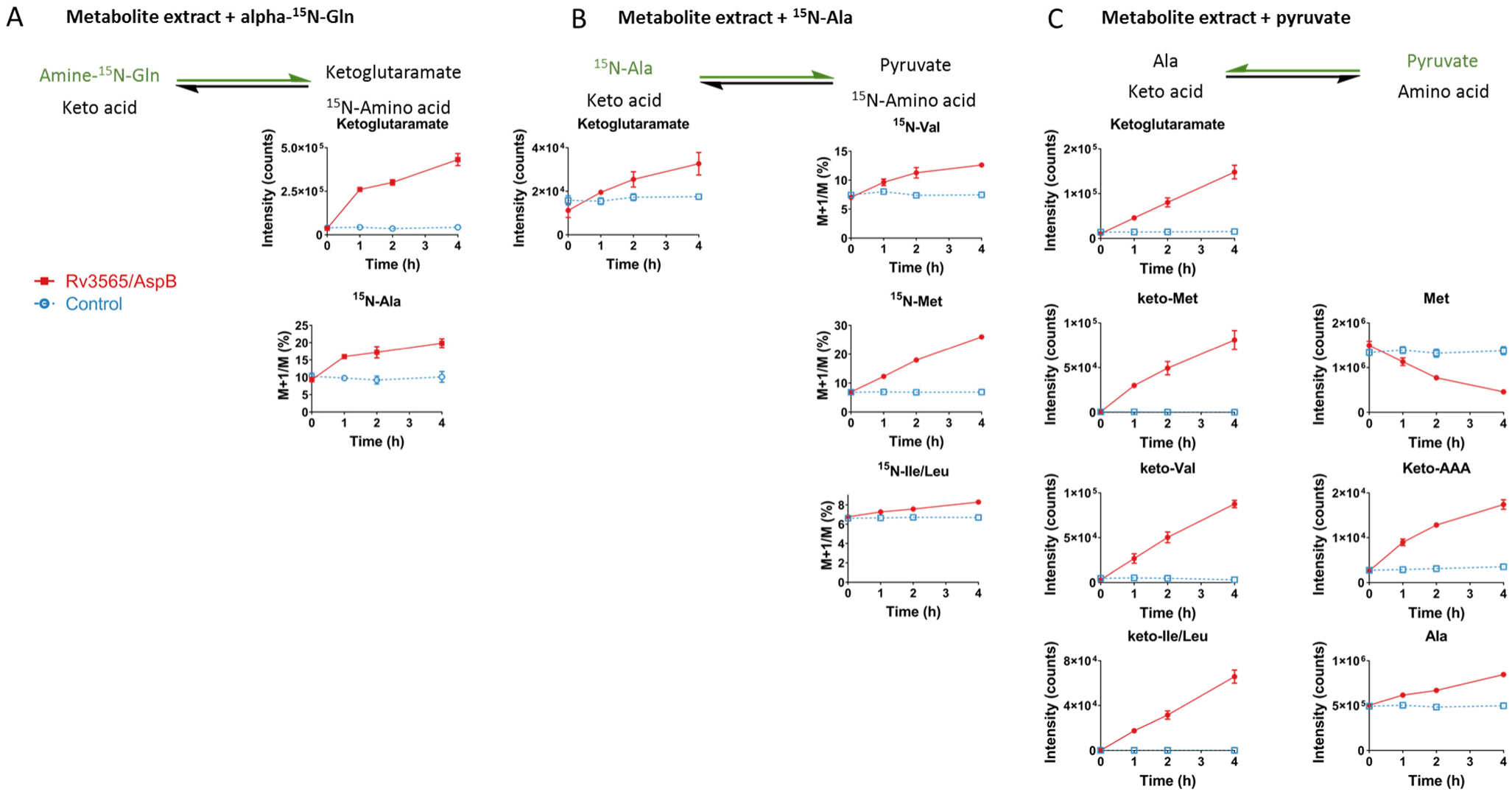
Rv3565/AspB functions as an aminotransferase. **A)** Activity-based metabolite profiling with Rv3565 in the presence of α-^15^N-Gln. Purified recombinant Rv3565 (10 µM; red line) or a buffer control, was incubated with a mycobacterial metabolite extract supplemented with 10 mM α-^15^N-Gln for 0, 1, 2 and 4 h at 37 °C, and analyzed using untargeted LC-MS. **B)** Activity-based metabolite profiling with Rv3565 in the presence of ^15^N-Ala and α-ketoglutarate. Same as A, but using a mycobacterial metabolite extract supplemented with 10 mM ^15^N-Ala. **C)** Activity-based metabolite profiling with Rv3565 in the presence of pyruvate. Same as A, but using a mycobacterial metabolite extract supplemented with 20 mM pyruvate. Colored arrows indicate the forced direction of the Rv3565-mediated reaction. Relative metabolite levels are represented as intensity, while ^15^N-labeling is presented as the ratio M+1/M, which was not corrected for naturally occuring isotopes. Keto-AAA, keto-Val, keto-Met, Keto-Ile/Leu: keto acids of the corresponding amino acids. Data are presented as mean +/- SD (n=3).

**Figure S13.**
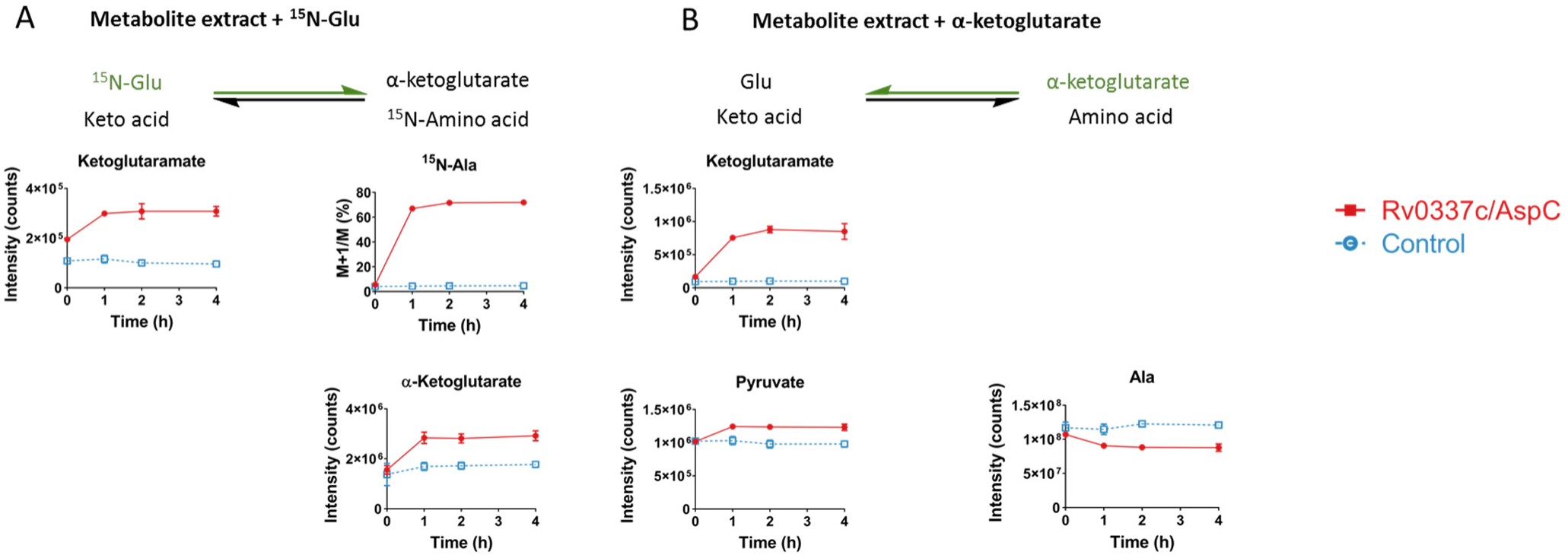
Rv0337c/AspC functions as an aminotransferase. **A)** Activity-based metabolite profiling with Rv0337c in the presence of α-^15^N-Gln. Purified recombinant Rv0337c (10 µM; red line) or a buffer control, was incubated with a mycobacterial metabolite extract supplemented with 10 mM α-^15^N-Gln for 0, 1, 2 and 4 h at 37 °C, and analyzed using untargeted LC-MS. **B)** Activity-based metabolite profiling with Rv0337c in the presence of α-ketoglutarate. Same as A, but using a mycobacterial metabolite extract supplemented with 20 mM α-ketoglutarate. Colored arrows indicate the forced direction of the Rv0337c-mediated reaction. Relative metabolite levels are represented as intensity, while ^15^N-labeling is presented as the ratio M+1/M, which was not corrected for naturally occuring isotopes. Data are presented as mean +/- SD (n=3).

**Figure S14.**
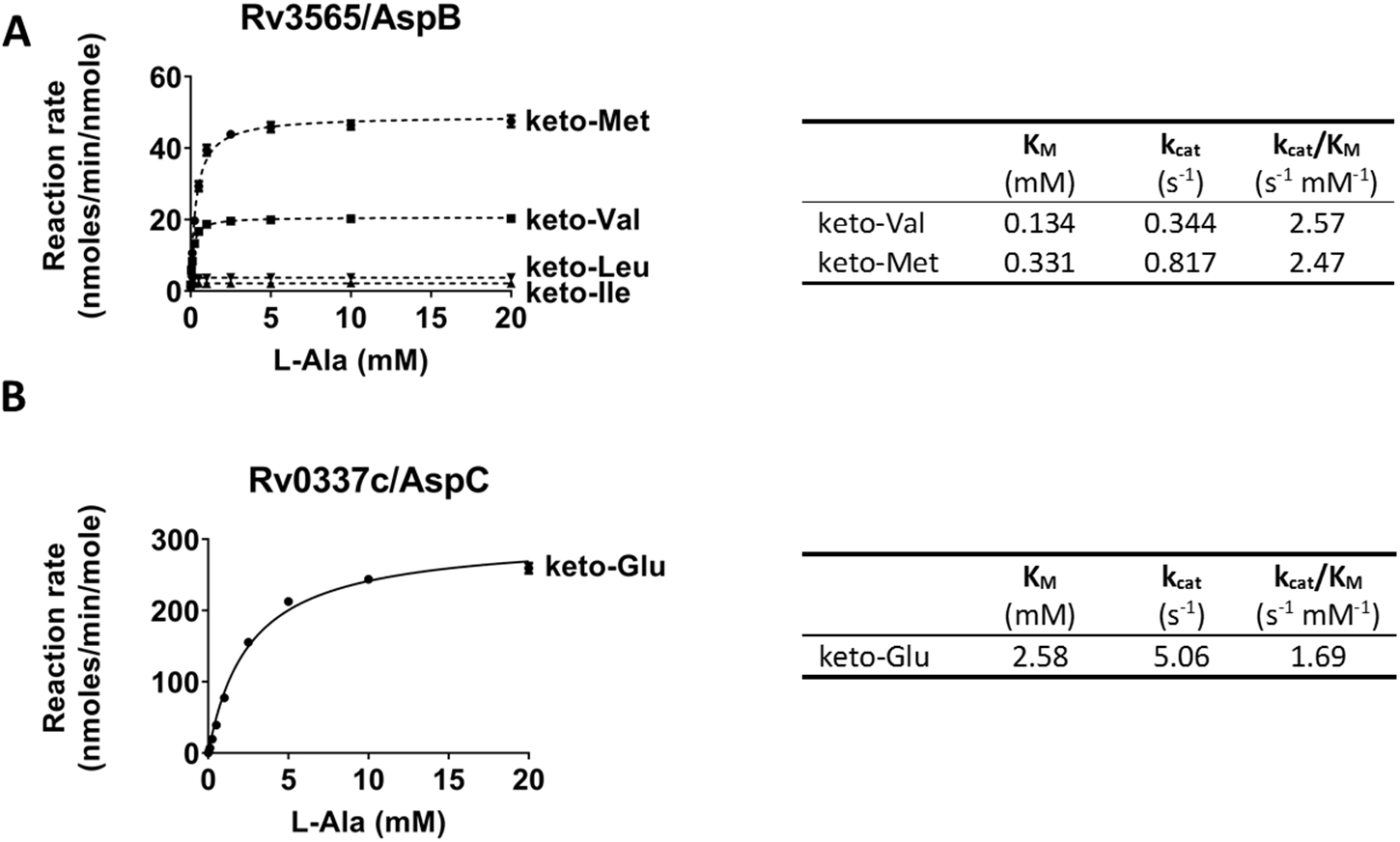
Rv3565/AspB functions as AvtA and Rv0337c/AspC as AlaT. **A)** Steady-state enzyme kinetics of Rv3565/AspB. Purified recombinant Rv3565 (1 µM) was incubated with 10 mM keto acids and increasing concentrations of alanine at 37 °C. Pyruvate formation was measured using a coupled reaction with lactate dehydrogenase. Measured velocities were fitted to Michaelis-Menten kinetics using Graphpad Prism 7 software. **B)** Steady-state enzyme kinetics of Rv0337c/AspC. Same as A, but for Rv0337c. Data are presented as mean +/- SD.

**Figure S15.**
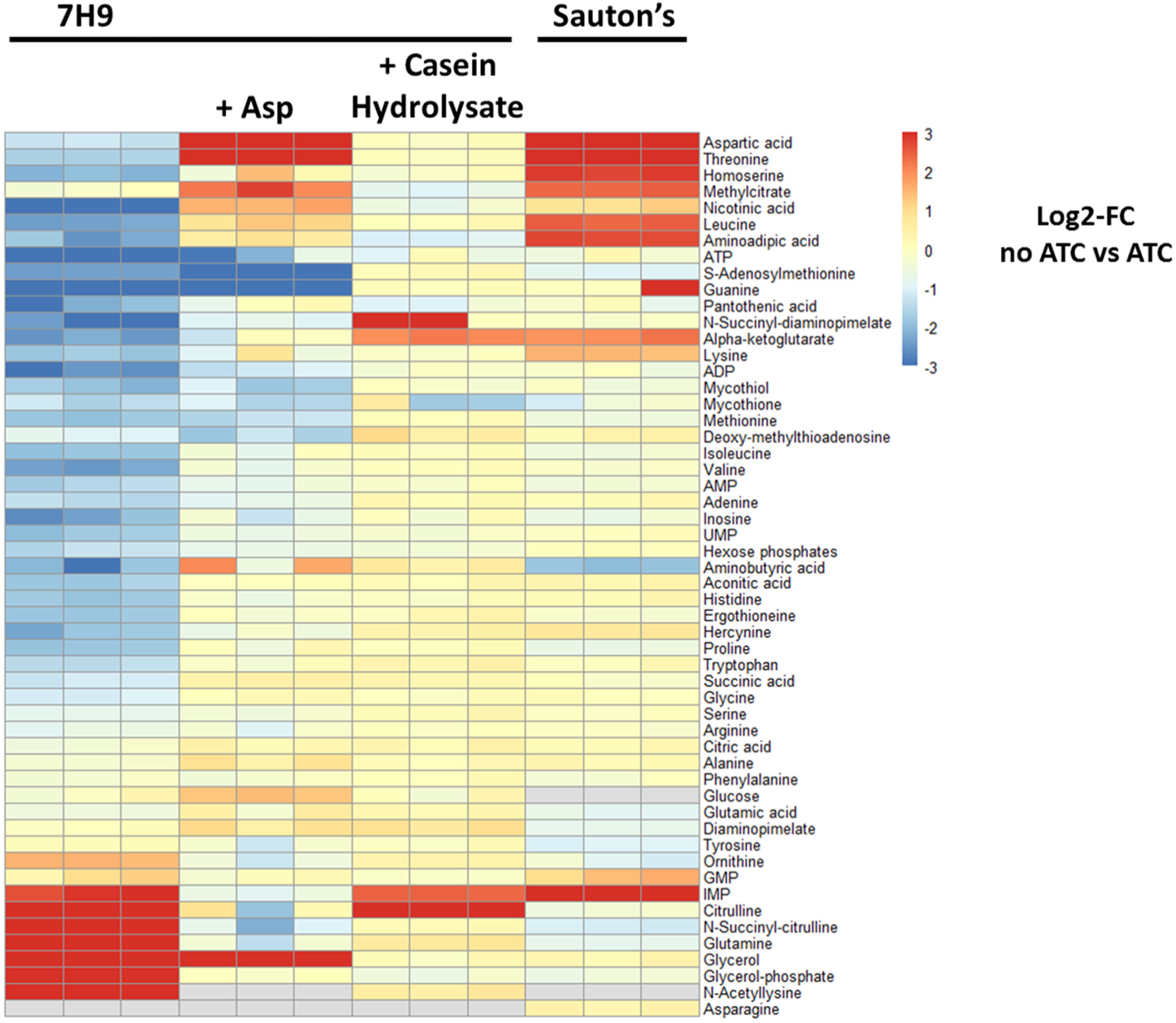
Rv3722c deficiency results in widespread metabolic changes. Heatmap showing the log2-fold change in metabolite levels in growth permissive and non-permissive culture media (Fig 1A, C, E and 7B). Rv3722c-TetON and Rv3722c-control, were cultured in 7H9, 7H9+3 mM aspartate, 7H9+ 1% casein hydrolysate (all with 0.04% tyloxapol), and Sauton’s minmal media, in the presence and absence of 500 ng/mL ATC. After 12 days, metabolite levels were obtained using high-resolution LC-MS. Colors represent log2-fold change of Rv3722c-deficient (no ATC) versus –proficient (ATC) Rv3772-TetON. Data are presented as mean ratio (n=3 vs n=3).

## References

1. Ellens, K. W. et al. Confronting the catalytic dark matter encoded by sequenced genomes. Nucleic Acids Res. 45, 11495–11514 (2017).

2. Galperin, M. Y. & Koonin, E. V. From complete genome sequence to ‘complete’ understanding? Trends Biotechnol. 28, 398–406 (2010).

3. Niehaus, T. D., Thamm, A. M. K., de Crécy-Lagard, V. & Hanson, A. D. Proteins of Unknown Biochemical Function: A Persistent Problem and a Roadmap to Help Overcome It. Plant Physiol. 169, 1436–1442 (2015).

4. Hanson, A. D., Pribat, A., Waller, J. C. & de Crécy-Lagard, V. ‘Unknown’ proteins and ‘orphan’ enzymes: the missing half of the engineering parts list--and how to find it. Biochem. J. 425, 1–11 (2009).

5. Chen, L. & Vitkup, D. Distribution of orphan metabolic activities. Trends Biotechnol. 25, 343–348 (2007).

6. WHO | Global tuberculosis report 2018. WHO Available at: http://www.who.int/tb/publications/global_report/en/. (Accessed: 9th August 2019)

7. Griffin, J. E. et al. High-resolution phenotypic profiling defines genes essential for mycobacterial growth and cholesterol catabolism. PLoS Pathog. 7, e1002251 (2011).

8. Sassetti, C. M., Boyd, D. H. & Rubin, E. J. Genes required for mycobacterial growth defined by high density mutagenesis. Mol. Microbiol. 48, 77–84 (2003).

9. El-Gebali, S. et al. The Pfam protein families database in 2019. Nucleic Acids Res. 47, D427–D432 (2019).

10. Mitchell, A. L., et al. InterPro in 2019: improving coverage, classification and access to protein sequence annotations. Nucleic Acids Res. 47, D351–D360 (2019).

11. Huerta-Cepas, J. et al. eggNOG 4.5: a hierarchical orthology framework with improved functional annotations for eukaryotic, prokaryotic and viral sequences. Nucleic Acids Res. 44, D286–D293 (2016).

12. Yang, Z., Zeng, X. & Tsui, S. K.-W. Investigating function roles of hypothetical proteins encoded by the Mycobacterium tuberculosis H37Rv genome. BMC Genomics 20, 394 (2019).

13. Ortega, C. et al. Systematic Survey of Serine Hydrolase Activity in Mycobacterium tuberculosis Defines Changes Associated with Persistence. Cell Chem. Biol. 23, 290–298 (2016).

14. Målen, H., Berven, F. S., Fladmark, K. E. & Wiker, H. G. Comprehensive analysis of exported proteins from Mycobacterium tuberculosis H37Rv. Proteomics 7, 1702–1718 (2007).

15. Penn, B. H. et al. An Mtb-Human Protein-Protein Interaction Map Identifies a Switch between Host Antiviral and Antibacterial Responses. Mol. Cell 71, 637–648.e5 (2018).

16. Ehrt, S., Schnappinger, D. & Rhee, K. Y. Metabolic principles of persistence and pathogenicity in Mycobacterium tuberculosis. Nat. Rev. Microbiol. 16, 496–507 (2018).

17. Gouzy, A., Poquet, Y. & Neyrolles, O. A central role for aspartate in Mycobacterium tuberculosis physiology and virulence. Front. Cell. Infect. Microbiol. 3, 68 (2013).

18. Gouzy, A. et al. Mycobacterium tuberculosis nitrogen assimilation and host colonization require aspartate. Nat. Chem. Biol. 9, 674–676 (2013).

19. Gouzy, A., Poquet, Y. & Neyrolles, O. Nitrogen metabolism in Mycobacterium tuberculosis physiology and virulence. Nat. Rev. Microbiol. 12, 729–737 (2014).

20. Johnson, E. O. et al. Large-scale chemical–genetics yields new M. tuberculosis inhibitor classes. Nature 1 (2019). doi:10.1038/s41586-019-1315-z

21. Kim, J.-H. et al. Protein inactivation in mycobacteria by controlled proteolysis and its application to deplete the beta subunit of RNA polymerase. Nucleic Acids Res. 39, 2210–2220 (2011).

22. de Carvalho, L. P. S. et al. Activity-based metabolomic profiling of enzymatic function: identification of Rv1248c as a mycobacterial 2-hydroxy-3-oxoadipate synthase. Chem. Biol. 17, 323–332 (2010).

23. Larrouy-Maumus, G. et al. Discovery of a glycerol 3-phosphate phosphatase reveals glycerophospholipid polar head recycling in Mycobacterium tuberculosis. Proc. Natl. Acad. Sci. 110, 11320–11325 (2013).

24. Saito, N. et al. Metabolomics approach for enzyme discovery. J. Proteome Res. 5, 1979–1987 (2006).

25. Sévin, D. C., Fuhrer, T., Zamboni, N. & Sauer, U. Nontargeted in vitro metabolomics for high-throughput identification of novel enzymes in Escherichia coli. Nat. Methods (2016). doi:10.1038/nmeth.4103

26. Shen, H. et al. The Human Knockout Gene CLYBL Connects Itaconate to Vitamin B12. Cell 171, 771–782.e11 (2017).

27. Cooper, A. J. L. & Kuhara, T. α-Ketoglutaramate: an overlooked metabolite of glutamine and a biomarker for hepatic encephalopathy and inborn errors of the urea cycle. Metab. Brain Dis. 29, 991–1006 (2014).

28. Guidetti, P., Amori, L., Sapko, M. T., Okuno, E. & Schwarcz, R. Mitochondrial aspartate aminotransferase: a third kynurenate-producing enzyme in the mammalian brain. J. Neurochem. 102, 103–111 (2007).

29. Han, Q., Fang, J. & Li, J. Kynurenine aminotransferase and glutamine transaminase K of Escherichia coli: identity with aspartate aminotransferase. Biochem. J. 360, 617–623 (2001).

30. Grishin, N. V., Phillips, M. A. & Goldsmith, E. J. Modeling of the spatial structure of eukaryotic ornithine decarboxylases. Protein Sci. Publ. Protein Soc. 4, 1291–1304 (1995).

31. Sigrist, C. J. A. et al. New and continuing developments at PROSITE. Nucleic Acids Res. 41, D344–347 (2013).

32. Son, H. F. & Kim, K.-J. Structural Insights into a Novel Class of Aspartate Aminotransferase from Corynebacterium glutamicum. PloS One 11, e0158402 (2016).

33. Dolzan, M. et al. Crystal structure and reactivity of YbdL from Escherichia coli identify a methionine aminotransferase function. FEBS Lett. 571, 141–146 (2004).

34. Mehta, P. K., Hale, T. I. & Christen, P. Aminotransferases: demonstration of homology and division into evolutionary subgroups. Eur. J. Biochem. 214, 549–561 (1993).

35. Malashkevich, V. N., Onuffer, J. J., Kirsch, J. F. & Jansonius, J. N. Alternating arginine-modulated substrate specificity in an engineered tyrosine aminotransferase. Nat. Struct. Biol. 2, 548–553 (1995).

36. Carroll, P., Pashley, C. A. & Parish, T. Functional analysis of GlnE, an essential adenylyl transferase in Mycobacterium tuberculosis. J. Bacteriol. 190, 4894–4902 (2008).

37. Yuan, J. et al. Metabolomics-driven quantitative analysis of ammonia assimilation in E. coli. Mol. Syst. Biol. 5, 302 (2009).

38. Catazaro, J., Caprez, A., Guru, A., Swanson, D. & Powers, R. Functional evolution of PLP-dependent enzymes based on active-site structural similarities. Proteins Struct. Funct. Bioinforma. 82, 2597–2608 (2014).

39. Muratore, K. E. et al. Molecular function prediction for a family exhibiting evolutionary tendencies toward substrate specificity swapping: recurrence of tyrosine aminotransferase activity in the Iα subfamily. Proteins 81, 1593–1609 (2013).

40. Onuffer, J. J. & Kirsch, J. F. Redesign of the substrate specificity of escherichia coli aspartate aminotransferase to that of escherichia coli tyrosine aminotransferase by homology modeling and site-directed mutagenesis. Protein Sci. 4, 1750–1757 (1995).

41. Cole, S. T. et al. Deciphering the biology of Mycobacterium tuberculosis from the complete genome sequence. Nature 393, 537–544 (1998).

42. Amorim Franco, T. M., Hegde, S. & Blanchard, J. S. Chemical Mechanism of the Branched-Chain Aminotransferase IlvE from Mycobacterium tuberculosis. Biochemistry 55, 6295–6303 (2016).

43. Bhor, V. M., Dev, S., Vasanthakumar, G. R. & Surolia, A. Spectral and kinetic characterization of 7,8-diaminopelargonic acid synthase from Mycobacterium tuberculosis. IUBMB Life 58, 225–233 (2006).

44. Nasir, N., Anant, A., Vyas, R. & Biswal, B. K. Crystal structures of Mycobacterium tuberculosis HspAT and ArAT reveal structural basis of their distinct substrate specificities. Sci. Rep. 6, 18880 (2016).

45. Holt, M. C. et al. Biochemical Characterization and Structure-Based Mutational Analysis Provide Insight into the Binding and Mechanism of Action of Novel Aspartate Aminotransferase Inhibitors. Biochemistry 57, 6604–6614 (2018).

46. Wrenger, C. et al. Specific inhibition of the aspartate aminotransferase of Plasmodium falciparum. J. Mol. Biol. 405, 956–971 (2011).

47. Gouzy, A. et al. Mycobacterium tuberculosis exploits asparagine to assimilate nitrogen and resist acid stress during infection. PLoS Pathog. 10, e1003928 (2014).

48. Tullius, M. V., Harth, G. & Horwitz, M. A. Glutamine synthetase GlnA1 is essential for growth of Mycobacterium tuberculosis in human THP-1 macrophages and guinea pigs. Infect. Immun. 71, 3927–3936 (2003).

49. Agapova, A. et al. Flexible nitrogen utilisation by the metabolic generalist pathogen Mycobacterium tuberculosis. eLife 8, (2019).

50. Jensen, R. A. & Calhoun, D. H. Intracellular roles of microbial aminotransferases: overlap enzymes across different biochemical pathways. Crit. Rev. Microbiol. 8, 229–266 (1981).

51. Lal, P. B., Schneider, B. L., Vu, K. & Reitzer, L. The redundant aminotransferases in lysine and arginine synthesis and the extent of aminotransferase redundancy in Escherichia coli. Mol. Microbiol. 94, 843–856 (2014).

52. Gelfand, D. H. & Steinberg, R. A. Escherichia coli mutants deficient in the aspartate and aromatic amino acid aminotransferases. J. Bacteriol. 130, 429–440 (1977).

53. Bennett, B. D. et al. Absolute metabolite concentrations and implied enzyme active site occupancy in Escherichia coli. Nat. Chem. Biol. 5, 593–599 (2009).

54. Soga, T. et al. Quantitative metabolome analysis using capillary electrophoresis mass spectrometry. J. Proteome Res. 2, 488–494 (2003).

55. Sugimoto, M. et al. MMMDB: Mouse Multiple Tissue Metabolome Database. Nucleic Acids Res. 40, D809–814 (2012).

56. Reitzer, L. Nitrogen assimilation and global regulation in Escherichia coli. Annu. Rev. Microbiol. 57, 155–176 (2003).

57. Reitzer, L. Biosynthesis of Glutamate, Aspartate, Asparagine, L-Alanine, and D-Alanine. EcoSal Plus 1, (2004).

58. Somashekar, B. S. et al. Metabolic Profiling of Lung Granuloma in Mycobacterium tuberculosis Infected Guinea Pigs: Ex vivo 1H Magic Angle Spinning NMR Studies. J. Proteome Res. 10, 4186–4195 (2011).

59. Bhattacharyya, N. et al. An Aspartate-Specific Solute-Binding Protein Regulates Protein Kinase G Activity To Control Glutamate Metabolism in Mycobacteria. mBio 9, e00931–18 (2018).

60. Rieck, B. et al. PknG senses amino acid availability to control metabolism and virulence of Mycobacterium tuberculosis. PLOS Pathog. 13, e1006399 (2017).

61. Murphy, K. C., Papavinasasundaram, K. & Sassetti, C. M. Mycobacterial recombineering. Methods Mol. Biol. Clifton NJ 1285, 177–199 (2015).

62. Nandakumar, M., Prosser, G. A., de Carvalho, L. P. S. & Rhee, K. Metabolomics of Mycobacterium tuberculosis. in Mycobacteria Protocols 105–115 (Humana Press, New York, NY, 2015). doi:10.1007/978-1-4939-2450-9_6

63. Pesek, J. J., Matyska, M. T., Loo, J. A., Fischer, S. M. & Sana, T. R. Analysis of hydrophilic metabolites in physiological fluids by HPLC-MS using a silica hydride-based stationary phase. J. Sep. Sci. 32, 2200–2208 (2009).

64. Pashley, C. A., Brown, A. C., Robertson, D. & Parish, T. Identification of the Mycobacterium tuberculosis GlnE promoter and its response to nitrogen availability. Microbiology 152, 2727–2734 (2006).

65. Rial, D. V. & Ceccarelli, E. A. Removal of DnaK contamination during fusion protein purifications. Protein Expr. Purif. 25, 503–507 (2002).

66. Jaisson, S., Veiga-da-Cunha, M. & Van Schaftingen, E. Molecular identification of omega-amidase, the enzyme that is functionally coupled with glutamine transaminases, as the putative tumor suppressor Nit2. Biochimie 91, 1066–1071 (2009).

67. Winn, M. D. et al. Overview of the CCP4 suite and current developments. Acta Crystallogr. D Biol. Crystallogr. 67, 235–242 (2011).

68. Vagin, A. & Teplyakov, A. Molecular replacement with MOLREP. Acta Crystallogr. D Biol. Crystallogr. 66, 22–25 (2010).

69. Kabsch, W. XDS. Acta Crystallogr. D Biol. Crystallogr. 66, 125–132 (2010).

70. Adams, P. D. et al. PHENIX: building new software for automated crystallographic structure determination. Acta Crystallogr. D Biol. Crystallogr. 58, 1948–1954 (2002).

71. Emsley, P., Lohkamp, B., Scott, W. G. & Cowtan, K. Features and development of Coot. Acta Crystallogr. D Biol. Crystallogr. 66, 486–501 (2010).

72. Liebschner, D. et al. Polder maps: improving OMIT maps by excluding bulk solvent. Acta Crystallogr. Sect. Struct. Biol. 73, 148–157 (2017).

73. Pettersen, E. F. et al. UCSF Chimera--a visualization system for exploratory research and analysis. J. Comput. Chem. 25, 1605–1612 (2004).

74. Mendler, K. et al. AnnoTree: visualization and exploration of a functionally annotated microbial tree of life. Nucleic Acids Res. 47, 4442–4448 (2019).

75. Letunic, I. & Bork, P. Interactive Tree Of Life (iTOL) v4: recent updates and new developments. Nucleic Acids Res. 47, W256–W259 (2019).

76. Zwart, P. H., Grosse-Kunstleve, R. W., Lebedev, A. A., Murshudov, G. N. & Adams, P. D. Surprises and pitfalls arising from (pseudo)symmetry. Acta Crystallogr. D Biol. Crystallogr. 64, 99–107 (2008).

77. Brooks, C. L. et al. Pseudo-symmetry and twinning in crystals of homologous antibody Fv fragments. Acta Crystallogr. D Biol. Crystallogr. 64, 1250–1258 (2008).

78. Gibrat, J. F., Madej, T. & Bryant, S. H. Surprising similarities in structure comparison. Curr. Opin. Struct. Biol. 6, 377–385 (1996).

79. Mehta, P. K., Hale, T. I. & Christen, P. Aminotransferases: demonstration of homology and division into evolutionary subgroups. Eur. J. Biochem. 214, 549–561 (1993).

80. Dolzan, M. et al. Crystal structure and reactivity of YbdL from Escherichia coli identify a methionine aminotransferase function. FEBS Lett. 571, 141–146 (2004).

